# Pitfalls and recommended strategies and metrics for suppressing motion artifacts in functional MRI

**DOI:** 10.1101/2021.09.18.460908

**Authors:** Vyom Raval, Kevin P. Nguyen, Marco Pinho, Richard B. Dewey, Madhukar Trivedi, Albert A. Montillo

**Affiliations:** University of Texas at Dallas, Richardson, Texas, USA; University of Texas Southwestern Medical Center, Dallas, Texas, USA

**Author notes:** University of Washington, Seattle, Washington, USA. Information sharing statement This work uses publicly available datasets, which are listed with hyperlinks in the Data Availability Statement. Code used for these analyses can be accessed through the hyperlink provided in the Methods.

**Keywords:** fMRI, motion artifact, pipeline, preprocessing

## Abstract

In resting-state functional magnetic resonance imaging (rs-fMRI), artefactual signals arising from subject motion can dwarf and obfuscate the neuronal activity signal. Typical motion correction approaches involve the generation of nuisance regressors, which are timeseries of non-brain signals regressed out of the fMRI timeseries to yield putatively artifact-free data. Recent work suggests that concatenating all regressors into a single regression model is more effective than the sequential application of individual regressors, which may reintroduce previously removed artifacts. This work compares 18 motion correction pipelines consisting of head motion, independent components analysis, and non-neuronal physiological signal regressors in sequential or concatenated combinations. The pipelines are evaluated on a dataset of cognitively normal individuals with repeat imaging and on datasets of studies of Autism Spectrum Disorder, Major Depressive Disorder, and Parkinson’s Disease. Extensive metrics of motion artifact removal are measured, including resting state network recovery, Quality Control-Functional Connectivity (QC-FC) correlation, distance-dependent artifact, network modularity, and test-retest reliability of multiple rs-fMRI analyses. The results reveal limitations in previously proposed metrics, including the QC-FC correlation and modularity quality, and identify more robust motion correction metrics. The results also reveal limitations in the concatenated regression approach, which is outperformed by the sequential regression approach in the test-retest reliability metrics. Finally, pipelines are recommended that perform well based on quantitative and qualitative comparisons across multiple datasets and robust metrics. These new insights and recommendations help address the need for effective motion artifact correction to reduce noise and confounds in rs-fMRI.

## Introduction

Resting-state fMRI (rs-fMRI), acquired in the absence of an active task, has become a popular method for studies of the brain’s intrinsic resting state networks (RSNs) and of functional connectivity (FC) between pairs of neuroanatomical regions (Fornito and Bullmore 2010; Smitha et al. 2017). However, analyses of rs-fMRI data are plagued by the presence of artifacts, which may seriously confound results or lead to spurious conclusions. Examples include physiological artifacts, where cardiac and respiratory activity cause signal fluctuations, and subject motion, which affects the image in several ways (Liu 2016). Motion within an axial image slice (i.e. anterior-posterior or left-right) changes which brain tissue is captured by each voxel, while motion across slices (i.e. superior-inferior) causes spin-history effects with complex effects on fMRI signal (Liu 2016; Friston et al. 1996). Motion-related signals can explain up to 30% of the total signal variance in fMRI, or about 1.6 times the signal change of neuronal activity (Bianciardi et al. 2009; Bright and Murphy 2015).

These motion artifacts may significantly confound FC measures in rs-fMRI (van Dijk et al. 2012). In one highly-cited example, the apparent relationship between age and brain network connectivity strength was found to be substantially inflated by motion, which itself was strongly correlated to subject age (Satterthwaite et al. 2012). Beyond FC, motion introduces confounds into most types of rs-fMRI analyses, including independent components analysis of RSNs, amplitude of low frequency fluctuation (ALFF), and fractional ALFF (fALFF) (Satterthwaite et al. 2012). Even fMRI acquired with subsecond temporal and spatial resolution below 3 mm, such as the Human Connectome Project, is susceptible to motion artifacts (Burgess et al. 2016). Consequently, there is a clear need for methods to correct for motion artifacts in rs-fMRI and to mitigate spurious findings from motion-contaminated analyses.

### Measurement of motion artifact contamination

Power et al. have defined metrics to quantify the efficacy of motion artifact correction methods (Power et al. 2015). These include the Quality Control-Functional Connectivity (QC-FC) metric, measuring the correlation between FC strength and subject motion (e.g. the mean framewise displacement), and the distance-dependent (QC-FC-dd) metric, which correlates the QC-FC for each inter-node brain connection with the connection’s anatomical length. Power et al. assert that successful motion artifact correction should reduce QC-FC correlation *and* distance-dependent artifact, as motion tends to preferentially inflate short-distance connectivity strength compared to longer-distance connectivity.

### Prospective motion artifact correction

Prospective methods for motion artifact correction seek to track subject motion in real-time and accordingly adjust scanner gradients, pulse frequency, and other imaging parameters. Unlike the retrospective methods addressed herein, prospective methods are able to directly correct for spin history effects and intra-volume distortions (Zaitsev et al. 2017; Huang et al. 2018; Maziero et al. 2020). However, the requisite equipment for these powerful methods, such as subject fiducials and an MR-compatible tracking system, lack widespread availability. It is also difficult to directly compare between methods, as there is no way to obtain a baseline uncorrected image or to apply multiple methods to the same image (Zaitsev et al. 2017). Consequently, this work focuses on retrospective methods, which are more widely applicable to any image from any scanner.

### Motion artifact regression methods

Retrospective motion artifact suppression methods assume the observed fMRI signal can be modeled with an additive model consisting of a linear combination of nuisance artifacts and the true brain signature. The artifacts are modeled as nuisance regressors, which form a design matrix ***X*** with dimensions of *t* timepoints ×*n* regressors. The artifacts are regressed out of the fMRI signal ***Y*** by solving for the coefficients ***β*** in the generalized linear model ***Y*** = ***Xβ***+ ***e***. This leaves the cleaned brain signal ***e***. Many methods have been proposed for the construction of nuisance regressors. Early approaches generated nuisance regressors from affine head motion parameters (HMP) (Power et al. 2012). HMP regression alone is insufficient to remove motion artifact (Satterthwaite et al. 2012; van Dijk et al. 2012; Power et al. 2012), since it does not account for longer-lasting spin history effects (Maknojia et al. 2019). Consequently, additional regressor types are needed, such as non-neuronal physiological signals measured from cerebrospinal fluid (CSF) and white matter (WM) (Power et al. 2012). Automated methods such as ICA-AROMA have been proposed to generate these regressors in a data-driven manner for more complete artifact removal (Pruim et al. 2015). Different combinations of regressors appear to be better at suppressing various types of artifacts, such as distance-dependent artifacts vs. global (whole-brain) artifacts (Burgess et al. 2016).

### Related work

There has been previous work to determine the optimal combinations and orderings of nuisance regressors (Ciric et al. 2017; Parkes et al. 2018). Ciric et al. compared multiple motion correction pipelines, consisting of various combinations of regression steps, using metrics such as QC-FC, QC-FC-dd, and network modularity (the ability of FC graphs to separate into networks). The comparison by Parkes et al. focused on the QC-FC, QC-FC-dd, and test-retest reliability metrics. While both studies evaluated multiple pipelines and metrics, there is yet room for further work on motion correction pipelines. *First*, these studies evaluated a limited set of metrics, specific to FC analyses: QC-FC, QC-FC-dd, test-retest reliability of FC, and network modularity. In this work, we also measure RSN identifiability in group independent component analysis (GICA) and seed-based connectivity (SBC) analyses. We also seek to address the hurdle of reproducibility in fMRI (Specht 2019). To investigate test-retest reliability across a variety of popularly analyzed rs-fMRI derivatives, we measure test-retest reliability not only in FC but also in GICA, SBC, fALFF, and regional homogeneity (ReHo) analyses. The *second* gap we address is the use of simultaneous nuisance regression. Recent work by Lindquist et al. equates nuisance regression steps to geometric projections onto an orthogonal subspace (Lindquist et al. 2019). While the result (i.e. ***e***, the cleaned data) of each regression step is orthogonal to its current regressors (**Figure 1a**), subsequent nuisance regressors are not necessarily orthogonal to previous steps (**Figure 1b**), and this may reintroduce previously removed artifacts (**Figure 1c**). Lindquist et al. recommend the use of a single regression step, where all nuisance regressors are concatenated into the same design matrix. Such an approach was not considered in the works of Ciric et al. and Parkes et al., so we will compare this *concatenated* regression with *sequential* regression in this work. *Finally*, our comparisons make use of multiple datasets, including data from neuropsychiatric diseases where greater subject motion may be encountered such as autism and Parkinson’s Disease. This breadth makes our results highly applicable to researchers applying rs-fMRI to the diseased brain.

**Figure 1.**
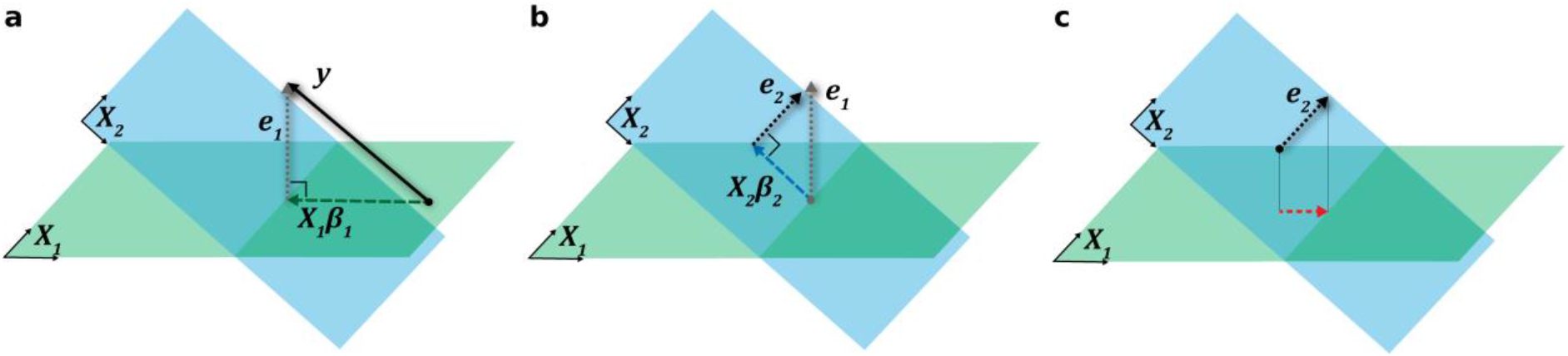
Sequential regression of data vector y on the hyperplanes ***X***_**1**_ (green) and ***X***_2_ (blue) defined by their associated nuisance regressors. **a**) ***y*** is projected onto ***X***_**1**_, resulting in ***X***_**1**_***β***_**1**_ which contains the portion of ***y*** explained by artifact ***X***_**1**_, while the residual ***e***_1_ is fully orthogonal to ***X***_**1**_ and contains the partially cleaned data. **b**) This data ***e***_1_ is next projected onto ***X***_**2**_, isolating the ***X***_2_ artifact ***X***_2_***β***_2_ and a new residual ***e***_2_ that is orthogonal to ***X***_**2**_. **c**) However, ***e***_2_ is no longer orthogonal to ***X***_**1**_. In general, ***e***_2_will have a component (red) that reintroduces the ***X***_**1**_artifact after the second regression if the two hyperplanes do not lie in the same subspace. This problem is a result of the fact that ***X***_**1**_ and ***X***_**2**_ are not regressed out together. If ***X***_**1**_ and ***X***_**2**_ were concatenated and regressed out together, the residual ***e*** would be orthogonal to both sets of regressors.

### Contributions

Through an extensive comparison of fMRI motion correction pipelines using several metrics and multiple datasets, this work makes the following contributions. 1) limitations in existing metrics are identified which may mislead comparisons of motion correction strategies and propose more reliable metrics. 2) evidence is presented indicating that the basic implementation of a concatenated regression does not work well in practice and propose several workarounds. 3) pipelines are recommended because they perform well across multiple sound metrics and are worth further investigation.

## Methods

### Materials

In this analysis, we used fMRI data from publicly available neuroimaging datasets, which were acquired with informed written consent and institutional review board approval at their respective institutions.

Data from the Human Connectome Project (HCP), one of the largest repositories of fMRI of healthy individuals, was used to compare the test-retest reliability (TRT) of fMRI processed by each of the motion correction pipelines. The S1200 release was used, which contains up to 4 resting-state scans for each subject (ages 22-35) acquired on the same day (van Essen et al. 2012). From this cohort, 50 cognitively healthy subjects were selected who had the greatest range in mean framewise displacement (mFD) across their scans and a minimum mFD below the cohort median (to ensure that at least one scan did not contain substantial motion artifact). The highest and lowest mFD images of each subject were included for analysis. See **Figure S1** for mFD distributions.

We additionally analyzed data from 3 datasets including subjects with psychiatric or neurodegenerative disease, where motion artifacts may be pervasive compared to cognitively normal cohorts. We investigated a cohort of young and adolescent autistic subjects aged 8-17 from the ABIDE I and ABIDE II databases (Di Martino et al. 2014). Three sites with the largest number of fMRI were included in this analysis: New York University (NYU) from ABIDE I (117 subjects), Kennedy Krieger Institute (KKI) from ABIDE II (151 subjects), and Georgetown University (GU) from ABIDE II (91 subjects). We also analyzed an adult psychiatric cohort, consisting of 270 subjects with major depressive disorder and 39 healthy controls aged 18-65 from the EMBARC study (Trivedi et al. 2016). Finally, we obtained data for 150 subjects aged 39-83 with Parkinson’s Disease, comprising an older cohort with a movement disorder, from the PPMI database^1^. In total, images from 868 subjects were included in this analysis.

Acquisition parameters and scanner information for these datasets can be found in **Table S1**. Demographics can be found in **Table S2**. The HCP images were considerably larger than the other datasets, with higher temporal resolution, spatial resolution, and acquisition time. To accommodate computer memory constraints and to make the TR of the HCP images more similar to that of the other datasets (2-2.5 s), we downsampled HCP images from the original TR = 0.72s to 2.88 s.

### Common Preprocessing Pipeline

All images were preprocessed with a common pipeline before performing motion correction. For fMRI preprocessing, we realigned volumes to correct for inter-volume head motion using the FSL MCFLIRT affine realignment tool (Jenkinson et al. 2002). We next performed skull-stripping by taking the intersection of brain masks generated by FSL BET and AFNI 3dAutomask (Smith 2002; Cox 1996). Afterwards, we spatially normalized images to the MNI152 EPI template using ANTs and normalized the intensity range to [0, 1000] (Avants et al. 2010). After the application of each specific motion correction pipeline, spatial smoothing was applied with a 4 mm FWHM kernel. An exception was made for pipelines using ICA-AROMA, where spatial smoothing was done *before* ICA-AROMA following developer recommendations (Pruim et al. 2015). For T1-weighted anatomical MRI, we performed skull-stripping with ROBEX and spatially normalized to the MNI152 T1-weighted template using ANTs (Iglesias et al. 2011; Avants et al. 2010). Finally, we segmented the anatomical MRI into gray matter, white matter, and CSF with FSL FAST (Zhang et al. 2001).

### Nuisance regressor generation

We generated four sets of nuisance regressors for each individual fMRI scan: 1) *Head motion parameters (HMP)* were computed during the affine volume realignment by FSL MCFLIRT. Along with the 6 original affine parameters, we included the first derivatives, squares, and squared derivatives for a total of 24 HMP regressors (Power et al. 2012; Satterthwaite et al. 2013). 2) *ICA-AROMA regressors* were identified with the ICA-AROMA package which decomposes the fMRI into independent components and classifies them as artifact or non-artifact signal (Pruim et al. 2015). 3) *Physiological artifact (Physio)* regressors included the mean white matter and CSF signals. Using the procedure recommended by (Power et al. 2018), we systematically eroded white matter and CSF masks from the anatomical image segmentation. The white matter mask was eroded for 5 cycles or until the next erosion would leave < 5 voxels, while the CSF mask was eroded twice. 4) *Frequency (Freq)* regressors were included to implement bandpass filtering at 0.008-0.08 Hz, using a set of sine and cosine timeseries generated with AFNI 1dBport (Cox 1996). Regression using these sinusoidal regressors is equivalent to bandpass filtering via Fast Fourier Transform yet allows testing simultaneous regression with the concatenated model.

### Pipelines evaluated

We utilized combinations of these regressors in 18 motion correction pipelines (**Table 1**). These included *sequential* pipelines in which nuisance regression was performed in successive steps and *concatenated* pipelines where all regressors were combined into one single design matrix. Within the sequential pipelines, we first combined HMP and AROMA in different orders (AROMA → HMP and HMP → AROMA), then appended Physio, and finally appended Freq. To create the concatenated pipelines, we started by combining HMP with either Physio or AROMA, then combined all three, and then added Freq. We also tested a [HMP, AROMA, Physio] → Freq pipeline where the high dimensionality Freq regressors were applied in a separate step. Due to the high dimensionality of the combined design matrix in the concatenated pipelines which may cause suboptimal regression model fitting, we tested a principal components regression (PCR) approach for the [HMP, AROMA, Physio] and [HMP, AROMA, Physio, Freq] pipelines. The concatenated design matrix was decomposed into the number of principal components that explain > 95% of the variance, and these components were used to form the new design matrix.

**Table 1.** The 18 motion correction pipelines compared in this study. The (→) indicates sequential regression steps while brackets ([x, y, z]) indicate concatenated regressors.

|  |
| --- |
| Baseline: no motion correction |
| Single algorithm: |
| HMP |
| AROMA |
| Physio |
| Freq |
| Sequential regression: |
| AROMA→HMP |
| HMP→AROMA |
| HMP→AROMA→Physio |
| AROMA→HMP→Physio |
| HMP→AROMA→Physio→Freq |
| AROMA→HMP→Physio→Freq |
| Concatenated regression |
| [HMP, Physio] |
| [HMP, AROMA] |
| [HMP, AROMA, Physio] |
| [HMP, AROMA, Physio, Freq] |
| [HMP, AROMA, Physio] → Freq |
| PCR [HMP, AROMA, Physio] |
| PCR [HMP, AROMA, Physio, Freq] |

We performed the regression using a Generalized Linear Model (GLM) via the regfilt tool in FSL. For pipelines including AROMA, we used the default “nonaggressive” approach, in which all AROMA regressors were included in the regression, including non-artifact regressors, and then the variance explained by the artifact regressors was subtracted from the data.

### Test-retest reliability

After processing the HCP dataset with each motion correction pipeline, we computed test-retest reliability (TRT) for 5 types of resting-state measurements between high- and low-motion images after motion correction. 1) Functional connectivity (FC) was computed by parcellating the images with the Gordon 333-region atlas and computing the pairwise correlations between mean regional signals (Gordon et al. 2016). We quantified TRT by computing the intraclass correlation (ICC) between the FC values from high- and low-motion images for each subject. 2) Regional homogeneity (ReHo) and 3) fractional amplitude of low frequency fluctuation (fALFF) were computed as additional measures of resting-state activity (Zang et al. 2004; Zou et al. 2008). We computed ReHo and fALFF using the C-PAC pipeline, which defines these measures at a specific band of 0.01-0.1 Hz (Craddock et al. 2013). Since this filter is standardized, we excluded the Freq regressors from each motion correction pipeline before computing these measures. We then computed mean regional values from ReHo and fALFF maps and computed ICC in the same manner as for FC. The final resting-state measurements were spatial maps of the default mode network (DMN), generated from each image using 4) group independent components analysis (GICA) and 5) seed-based connectivity (SBC). The DMN was chosen since it has been shown to be the most reliable brain network in resting-state fMRI (Franco et al. 2009; Beckmann et al. 2005; Greicius et al. 2003). After generating subject-specific DMN maps using GICA and SBC, spatial correspondence between the DMN recovered from each subject’s high motion image vs. their low motion image was measured using the Dice similarity coefficient (Thomason et al. 2011).

GICA was conducted with the GIFT tool to decompose fMRI into RSNs (Calhoun et al. 2001). GICA outputs a group-level mean decomposition as well as back-reconstructed subject-level decompositions, and the following optimization of parameters was done at the subject-level. The number of independent components (ICs) was optimized through an unbiased grid search to recover an IC with the highest spatial correspondence to a DMN template (Thomason et al. 2011). We searched for the optimal number of components (*k*) over the range of 12-24. Each output IC was then z-normalized and thresholded to create a binarized spatial map. We also optimized the threshold (*t*) via grid search over the range 0.2-2.0 in steps of 0.1. We selected the *k* x *t* combination which produced the maximum Dice similarity between each IC and the DMN template. This optimization was done separately for each pipeline to estimate each pipeline’s ability to recover the DMN. Seed-based connectivity (SBC) maps were generated using a 6 mm radius seed at the right posterior cingulate cortex at MNI coordinate (4, -54, 26) (Franco et al. 2009). We computed connectivity between each voxel and the seed using Pearson’s correlation, then converted correlation values to Z-scores using Fisher’s transform and applied a threshold at Z-score of 0.4.

### Quality control metrics

After processing the ABIDE, EMBARC, and PPMI datasets with each pipeline, we computed the metrics shown in **Figure 2**. First, we computed FC matrices from each image using the Gordon 333-region atlas. We also computed the mean framewise displacement (mFD) of each image using the HMP timeseries. For a subject’s image containing a timeseries of *n* volumes:

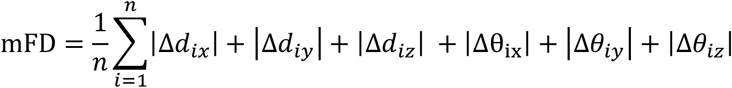

**Figure 2.**
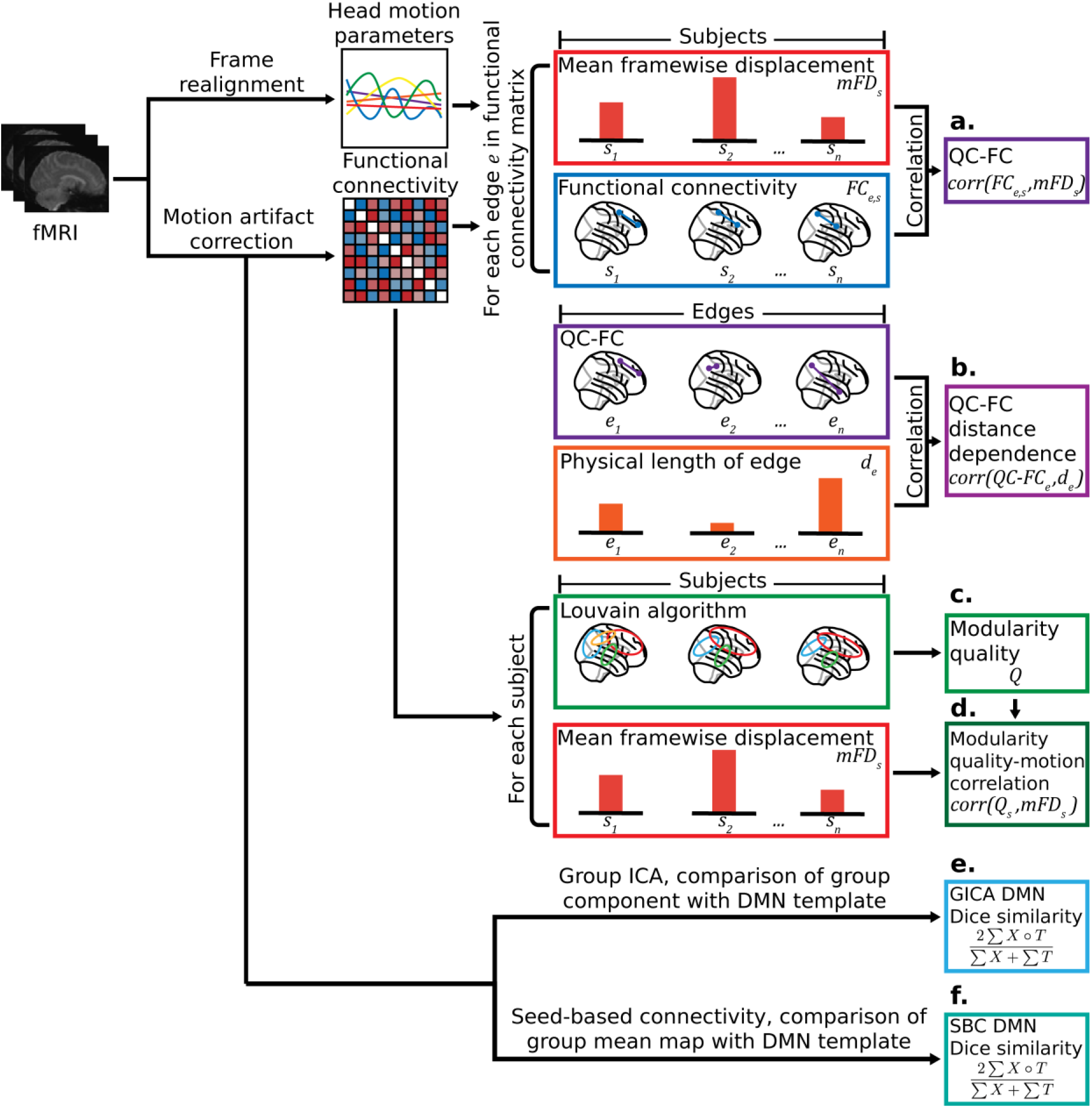
Computation steps for the six QC metrics (a)-(f). The frames of each subject’s raw fMRI are realigned which gives the head motion parameters (three rotation and three translation parameters). The image is then processed separately with each of the motion correction pipelines. The functional connectivity matrix is used to compute (a) QC-FC, (b) QC-FC distance dependence, (c) modularity quality, and (d) modularity quality-motion correlation. The motion-corrected images are also used to compute (e) the Dice Similarity coefficient (DSC) between a default mode network (DMN) template from an external dataset and the group ICA (GICA) DMN component, and (f) the DSC between the template DMN and each subject’s seed-based connectivity (SBC) map of the DMN.

where Δ*d*_*ix*_, Δ*d*_*iy*_, Δ*d*_*iz*_ are change in translation between volume *i* and the preceding volume *i* − 1 in the *x, y, z* dimensions (e.g. Δ*d*_*ix*_ = *d*_*ix*_ − *d*_(*i*−1)*x*_) and Δ*θ*_*ix*_, Δ*θ*_*iy*_, Δ*θ*_*iz*_ are the change in rotation (Power et al. 2012). Rotational displacements were converted from radians to millimeters assuming a mean brain radius of 50 mm.

We computed the *QC-FC correlation* and *QC-FC distance dependence* metrics as defined in (Power et al. 2015; Ciric et al. 2017; Parkes et al. 2018). For each edge in the FC matrix, QC-FC correlation (**Figure 2a**) is computed as the Pearson’s correlation between the mFD of each subject *s* and their FC strength for that edge

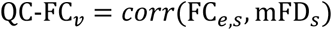

for each edge *e* in the FC matrix. The Gordon 333-region atlas yields an FC matrix containing 55,278 edges in its upper triangle. QC-FC values closer to 0 indicate that FC is less correlated with motion. The second metric, QC-FC distance dependence (**Figure 2b**), is Spearman’s rank correlation coefficient between the physical length of an edge in the brain *d*_*e*_ and its QC-FC:

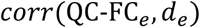

A QC-FC distance dependence closer to 0 indicates less distance-dependent motion contamination of FC. Spearman’s rank correlation was chosen here over Pearson’s correlation to better account for non-linear associations (Parkes et al. 2018). From the FC matrices, we also computed network modularity *Q* (**Figure 2c**) as described in (Ciric et al. 2017; Satterthwaite et al. 2012; Satterthwaite et al. 2013). The metric *Q*, bounded in the range [-1/2, 1], measures the difference between the number of within-network edges and the number of between-network edges (Newman 2006). The networks were detected from each FC matrix using the Louvain algorithm, a greedy optimization method that finds the graph communities that maximize *Q* (Blondel et al. 2008). From this modularity metric, we derived the modularity quality-motion correlation metric (**Figure 2d**), which is the Spearman’s rank correlation between a subject’s *Q*_*s*_ and their mFD:

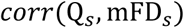

If motion artifacts do not contaminate FC matrices or adversely affect the community partitioning of the Louvain algorithm, which in turn affects modularity, there should be no correlation between modularity *Q* and motion mFD. For all correlation metrics, absolute values were used to index the strength of association and the direction of association was not considered.

Additional metrics were derived from resting state network (RSN) analyses of the processed images. As with the HCP dataset, we used GICA and SBC to generate DMN spatial maps. We created group-level maps using GICA, and for SBC, we computed the mean DMN map across all subjects. The GICA and SBC-based maps were binarized and the Dice similarity coefficient was computed with respect to a DMN template (Thomason et al. 2011)^2^. The binarization threshold was again optimized using a grid search to maximize Dice similarity and to provide the most optimistic estimate of pipeline performance, however this time the optimization was performed at the group-level (yielding one threshold per pipeline).

Code used to compute the metrics for these analyses has been made publicly available at https://gitfront.io/r/DeepLearningForPrecisionHealthLab/8c101f7218a58aabfdb14a7187eb0ad59b96021c/fMRI-Motion-Artifact-Suppression/.

## Results

### Test-retest reliability

Test-retest reliability measurements between paired, same-day images of the same subject in HCP showed substantial differences across pipelines. For brevity, we present here the results for the 5 pipelines that included all regressor types (HMP, Physio, AROMA, and Freq) because a typical fMRI analysis requires the removal of each of these artifacts. Full results for all 18 pipelines can be found in the Supplement, **Figures S2-S5**. Measurements of FC demonstrated the highest intraclass correlation (ICC) with the fully sequential HMP→AROMA→Physio→Freq pipeline (**Figure 3a**), which significantly outperformed the other 4 pipelines (paired t-test, *p* < 0.001). The other sequential pipeline AROMA→HMP → Physio→Freq ranked second, followed by the 3 concatenated pipelines. Notably, the fully concatenated [HMP, AROMA, Physio, Freq] pipeline showed worse FC ICC than the baseline images before any artifact correction. For ReHo and fALFF, the sequential pipelines also yielded significantly better ICC (*p* < 0.001) than the concatenated pipelines (**Figure 3b**,**c**). Next, the GICA-based, subject-specific DMN spatial maps demonstrated the best TRT (Dice similarity coefficient between paired images of the same subject) after processing with the PCR[HMP, AROMA, Physio, Freq] pipeline, followed by the sequential HMP→AROMA→Physio→Freq pipeline (**Figure 3d**). Comparisons of *group*-level DMN spatial maps in **Figure 4** showed similar TRT (Dice similarity between the low motion and high motion maps) across the pipelines, with the [HMP, AROMA, Physio] →Freq pipeline ranking first. Dice similarity between these DMN maps and the Thomason et al. 2011 template was also comparable across pipelines (**Figure S6**). However, qualitative differences appeared (**Figure 4**), such as the [HMP, AROMA, Physio] →Freq and [HMP, AROMA, Physio, Freq] pipelines failing to recover a significant prefrontal region of the DMN in either the low or high motion images. On the other hand, the SBC-based DMN spatial map showed the best TRT with the HMP→AROMA→Physio→Freq pipeline, with the sequential pipelines outperforming the concatenated pipelines (**Figure 3e**). SBC-based DMN TRT with the fully concatenated [HMP, AROMA, Physio, Freq] pipeline was worse than in the baseline images. Example DMN spatial maps for a representative subject are compared qualitatively in supplemental **Figure S7**. The baseline images produce noisy DMN maps that fail to recover the characteristic regions of the DMN, including the medial prefrontal cortex, posterior cingulate cortex, posterior parietal lobule, and hippocampus (Franco et al. 2009; Beckmann et al. 2005; Greicius et al. 2003). Meanwhile, these regions are recovered in the DMN map from the HMP→AROMA→Physio→Freq pipeline. The [HMP, AROMA, Physio] →Freq pipeline also recovered these regions, though its DMN map contains more noise. The [HMP, AROMA, Physio, Freq] pipeline, which had the worst TRT, contained the most noise.

**Figure 3.**
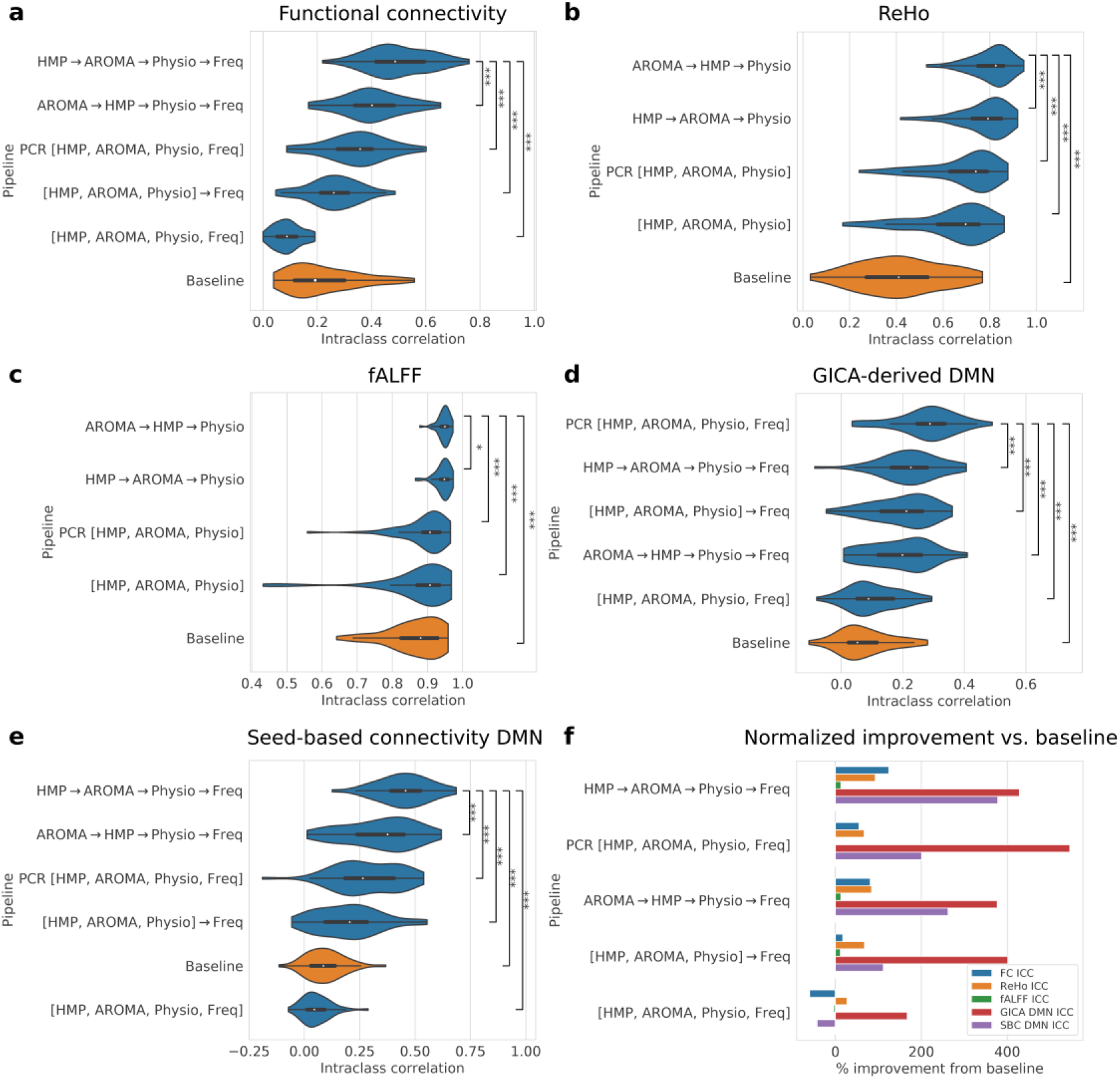
Comparison of pipeline test-retest reliability using resting-state fMRI connectivity measures. In this comparison, paired high- and low-motion images acquired in the same subject on the same day from the Human Connectome Project were used. The comparison is limited to the pipelines that include all 4 nuisance regressor types: head motion parameters (HMP), AROMA, physiological regressors, and frequency. Baseline is the uncorrected data. **a**) Processed images were parcellated with the Gordon 333-ROI atlas and the functional connectivity (FC) matrix was computed. The intraclass correlation (ICC) was computed between the high- and low-motion FC matrices of each subject and the distribution of subjects is shown. ICC was also computed for **b**) Regional homogeneity (ReHo) and **c**) fractional amplitude of low frequency fluctuations (fALFF). Here, frequency filtering was performed in a separate step from motion correction since the computation of ReHo and fALFF require their own bandpass filters. Spatial maps of the DMN were computed using **d**) group ICA and **e**) seed-based connectivity with a posterior cingulate cortex seed, and ICC was computed between high- and low-motion DMN maps of each subject. Pipelines significantly different from the top-ranking pipeline are annotated: * p < 0.05, ** p < 0.01, *** p < 0.001. In **f**), these 5 metrics are presented as normalized percentage improvement vs. baseline. Pipelines are ordered by mean overall percentage improvement.

**Figure 4.**
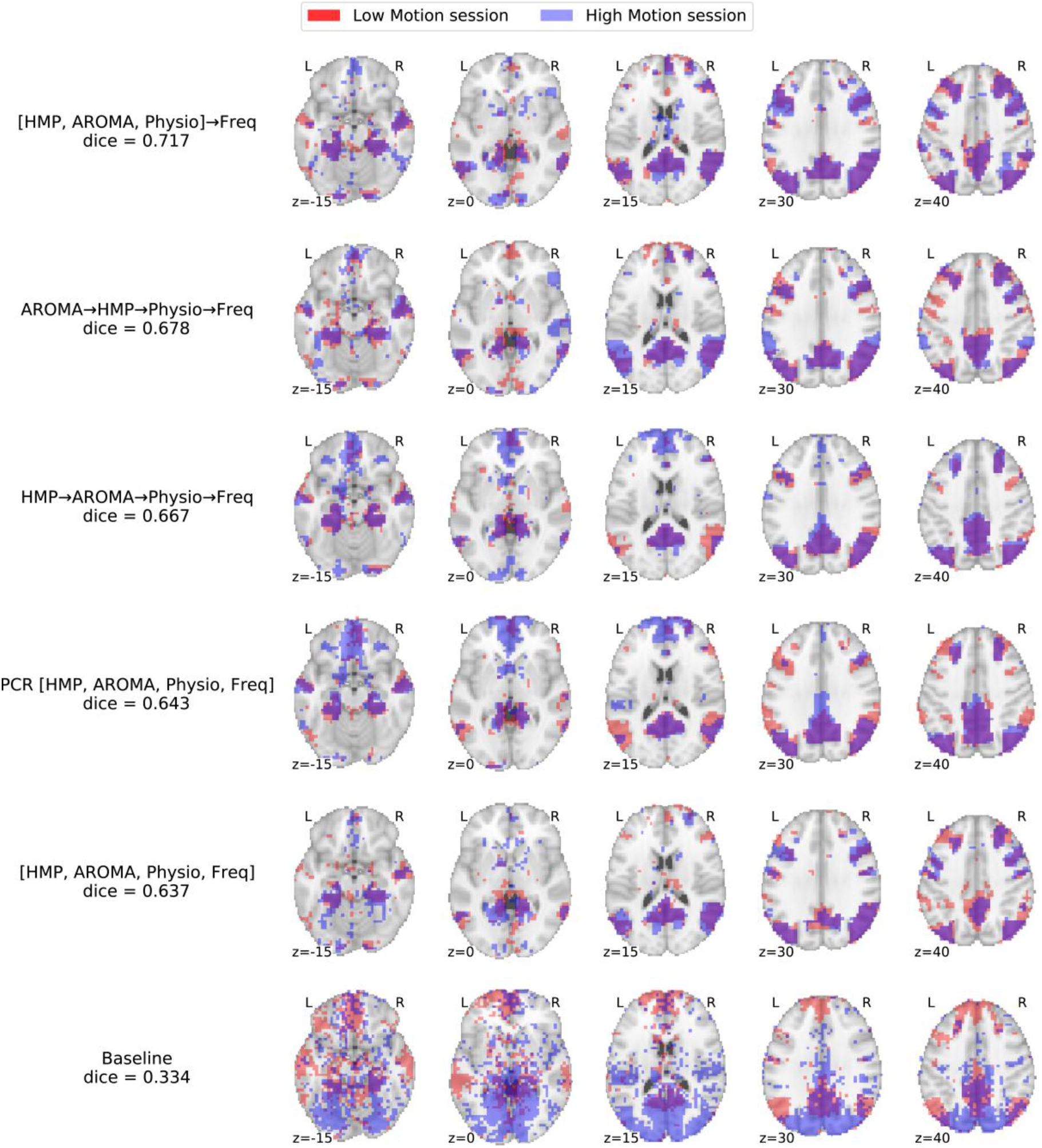
Graphical comparison of pipeline test-retest reliability of group-level default mode network (DMN) maps computed with group ICA (GICA) in the HCP dataset. GICA was performed separately on the 50 low motion images and 50 high motion images selected from the HCP dataset, and the independent component corresponding to the DMN was chosen using the described template-matching grid search. Dice similarity coefficient between the low motion and high motion session DMN spatial maps was computed as a measure of test-retest reliability for each pipeline. Shown here are the overlaid maps, with Low Motion map in red, High Motion map in blue, and overlapping areas in purple. Pipelines containing all 4 sets of regressors are shown in decreasing order of Dice similarity coefficient. Results from the baseline (no motion correction) pipeline are displayed on the bottom to illustrate the noisy GICA result before motion correction.

In summary, we found that the fully sequential pipelines, i.e. HMP→AROMA→Physio→Freq and AROMA→HMP→Physio→Freq, ranked first or second in all 5 resting-state, within-subject TRT measures. The fully concatenated [HMP, AROMA, Physio, Freq] pipeline, however, performed the poorest in all 5 measures.

### Default mode network recovery in disease datasets

Using the ABIDE, EMBARC, and PPMI datasets, we further compared the pipelines in their ability to recover the DMN in the presence of neurological and psychiatric disorders. As above, we present the subset of results from the pipelines containing all 4 types of nuisance regressors. Full results for all 18 pipelines tested can be found in **Figures S8 and S9**. After computing a group-level DMN map using GICA, 4 of the 5 compared pipelines showed comparable Dice similarity with the DMN template, with the AROMA→HMP→Physio→Freq pipeline ranking first (**Figure 5a**). The finding was similar for the SBC-based DMN maps, with 4 of the 5 pipelines scoring similarly and the PCR[HMP, AROMA, Physio, Freq] pipeline ranking first (**Figure 5b**). In both the GICA and SBC analyses, the fully concatenated pipeline [HMP, AROMA, Physio, Freq] exhibited the worst performance, with lower Dice similarity with the template compared to even the baseline.

**Figure 5.**
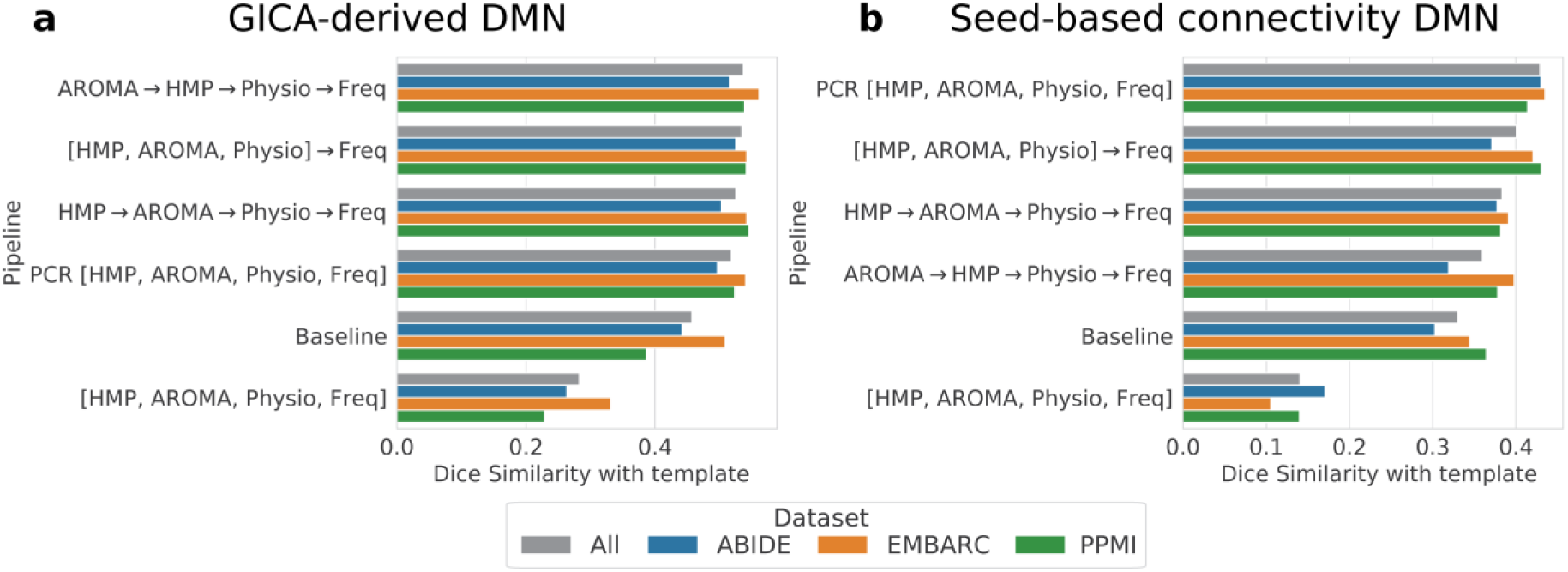
Default mode network (DMN) recovery by each pipeline in datasets containing neurological and psychiatric disorders. a) GICA and b) SBC were used to create group-level DMN maps for 3 neuropsychiatric disorder datasets: ABIDE (Autism Spectrum Disorder), EMBARC (Major Depressive Disorder), and PPMI (Parkinson’s Disease). Dice similarity was computed between these DMN maps and a standard template. Higher Dice similarity indicates better recovery of the canonical DMN. Pipelines are sorted in descending order of the average Dice similarity across the 3 datasets weighted by number of subjects (“All” score in gray).

A qualitative comparison of GICA DMN maps across pipelines is shown in **Figure 6**, for KKI site of the ABIDE dataset. Findings were similar across the rest of the ABIDE, EMBARC, and PPMI datasets. Of note, the baseline images failed to recover the hippocampal region of the DMN. The PCR[HMP, AROMA, Physio, Freq] pipeline did recover the hippocampus but missed the parietal lobe regions. The HMP→ AROMA→Physio→Freq, AROMA→HMP→Physio→Freq, and [HMP, AROMA, Physio] →Freq pipelines were each capable of recovering the key DMN regions to varying degrees. Consistent with the quantitative observation in **Figure 5**, the [HMP, AROMA, Physio, Freq] pipeline failed to recover any meaningful structures of the DMN (**Figure 6**).

**Figure 6.**
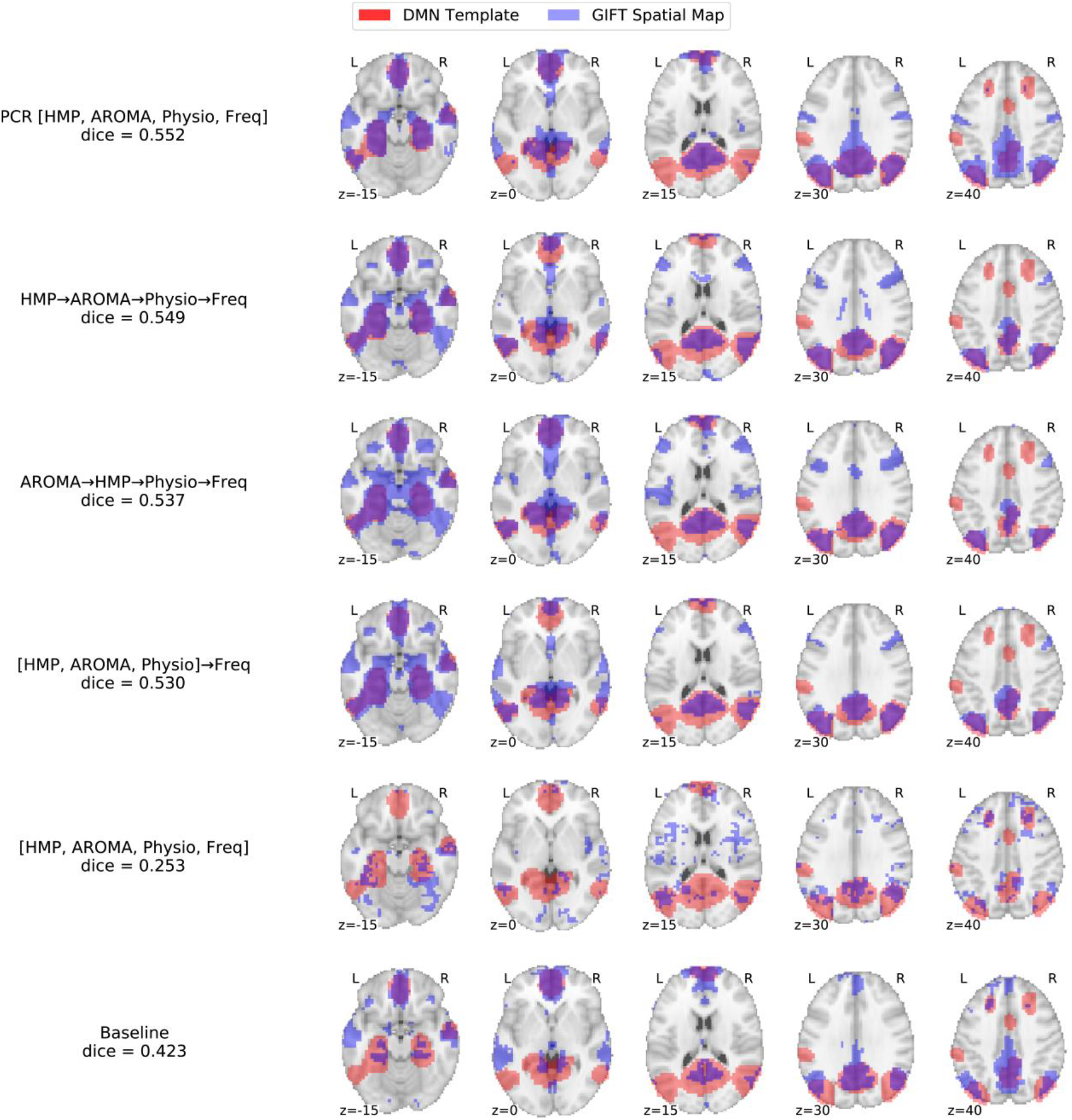
Graphical comparison of the pipelines’ test-retest reliability of group-level default mode network (DMN) maps computed with group ICA (GICA) in the dataset of subjects with Autism Spectrum Disorder. GICA using GIFT was computed on the 151 subjects (8-17 years old) from the ABIDE2KKI dataset, and the IC corresponding to the DMN was chosen using the described template-matching optimization method. Shown here are the template DMN in red and GIFT extracted DMN in blue, with overlapping areas in purple. Pipelines with all 4 sets of regressors are shown in decreasing order of dice. Baseline pipeline is displayed on the bottom, indicating that GIFT is unable to recover the hippocampal areas without motion correction. Notably, GIFT is not able to recover a meaningful DMN with the [HMP, AROMA, Physio, Freq] pipeline, which was also the case for all datasets except ABIDE2GU, indicating that the pipeline may remove DMN signal in many cases.

### Functional connectivity metrics in disease datasets

As a primary purpose for FC measures is the diagnosis and prognosis of neurologic disease, we did an additional exploratory analysis of FC-based motion correction metrics in the disease datasets specifically. The FC metrics evaluated in the ABIDE, EMBARC, and PPMI datasets appeared to support a different ranking of pipelines. This was the case for the 5 pipelines containing all 4 types of nuisance regressors (**Figure 7**), and for the results for all 18 pipelines tested which are shown in **Figures S10-S13**. All pipelines were able to improve QC-FC correlation and QC-FC-dd from baseline, indicating removal of motion confound and distance-dependent artifact from FC matrices. All pipelines were also able to improve modularity quality (*Q*), indicating better graph partitioning of the FC matrices. For the modularity quality-motion correlation metric, all pipelines except [HMP, AROMA, Physio, Freq] were able to achieve improvements over baseline. Further, for QC-FC and modularity quality, the [HMP, AROMA, Physio, Freq] pipeline ranked first among the pipelines, despite having performed poorly in the DMN recovery metrics.

**Figure 7.**
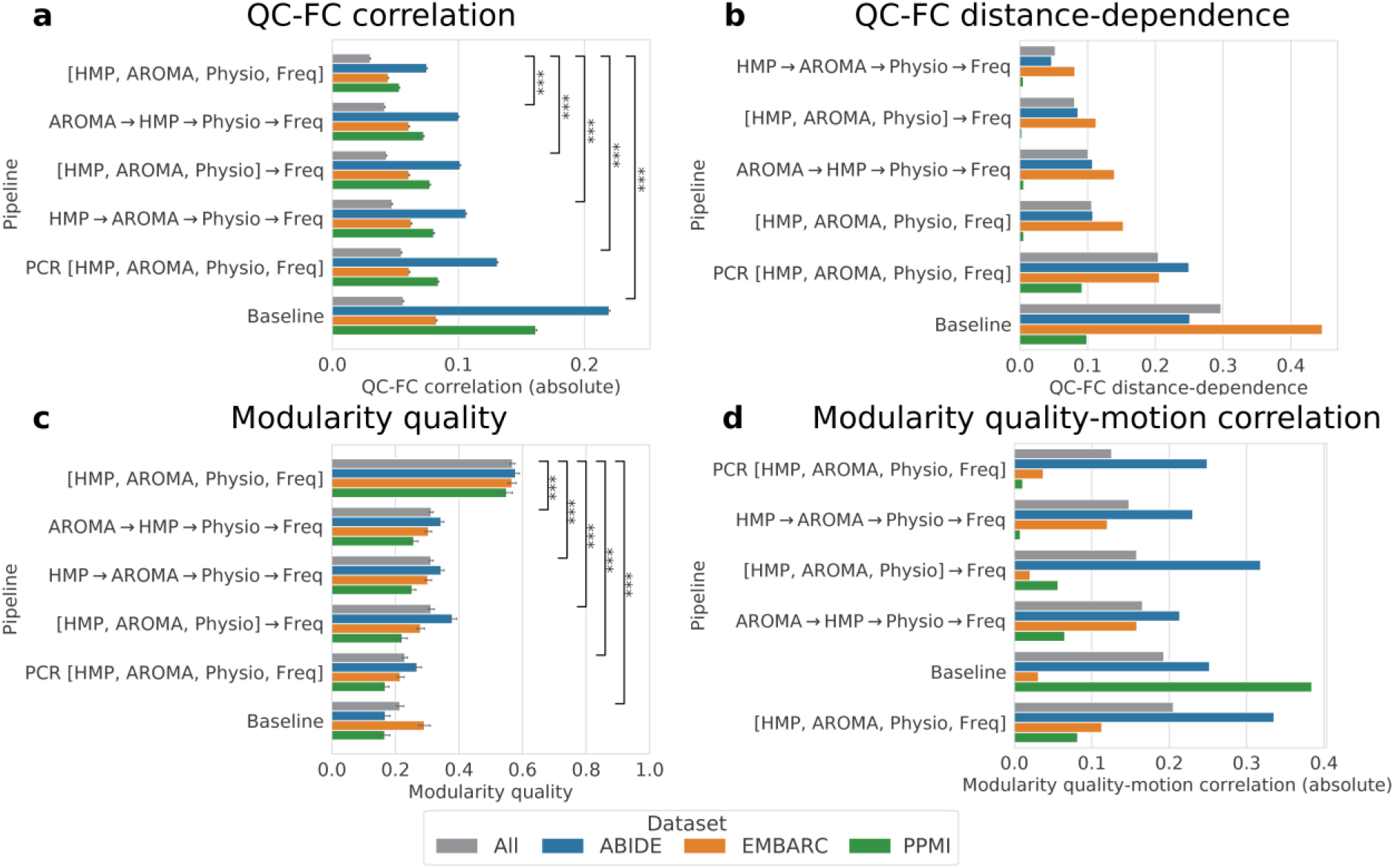
Additional quality metrics compared across pipelines for the 3 neuropsychiatric disease datasets: ABIDE (Autism Spectrum Disorder), EMBARC (Major Depressive Disorder), and PPMI (Parkinson’s Disease). The pipelines are sorted from best (top) to worst (bottom) by the “All” score (in gray), which is the average across the 3 datasets weighted by number of subjects. Metrics computed included a) QC-FC correlation, b) QC-FC distance dependence, c) modularity quality, and d) modularity quality-motion correlation. See the text for a discussion on the validity of these metrics. For a) QC-FC correlation which is a distribution over FC edges and c) modularity quality which is a distribution over subjects, 95% confidence intervals are included as error bars. Significant differences in “All” score for each pipeline vs. baseline are indicated with asterisks: *** p < 0.001.

These apparent contradictions led us to more closely test the validity of several widely used quality metrics.

### Closer examination of the QC-FC correlation metric

We further investigated the discrepancy between the DMN metrics and the FC metrics, specifically the QC-FC correlation. **Figure 8** compares how the GICA DMN Dice similarity and QC-FC correlation associate with different measures of DMN connectivity. The ground truth for DMN Dice similarity was the RSN labels of the Gordon-333 atlas. Overall *DMN connectivity strength*, computed from all FC matrices as the mean intra-DMN functional connectivity, was positively correlated with GICA DMN Dice similarity and negatively correlated with QC-FC correlation across the 18 pipelines. Likewise, overall DMN homogeneity, computed as the negative mean of per-subject intra-DMN connectivity standard deviations, was also positively correlated with GICA DMN Dice similarity and negatively correlated with QC-FC correlation. Consequently, *higher* (*better)* QC-FC was associated with *lower (poorer)* DMN connectivity and homogeneity.

**Figure 8.**
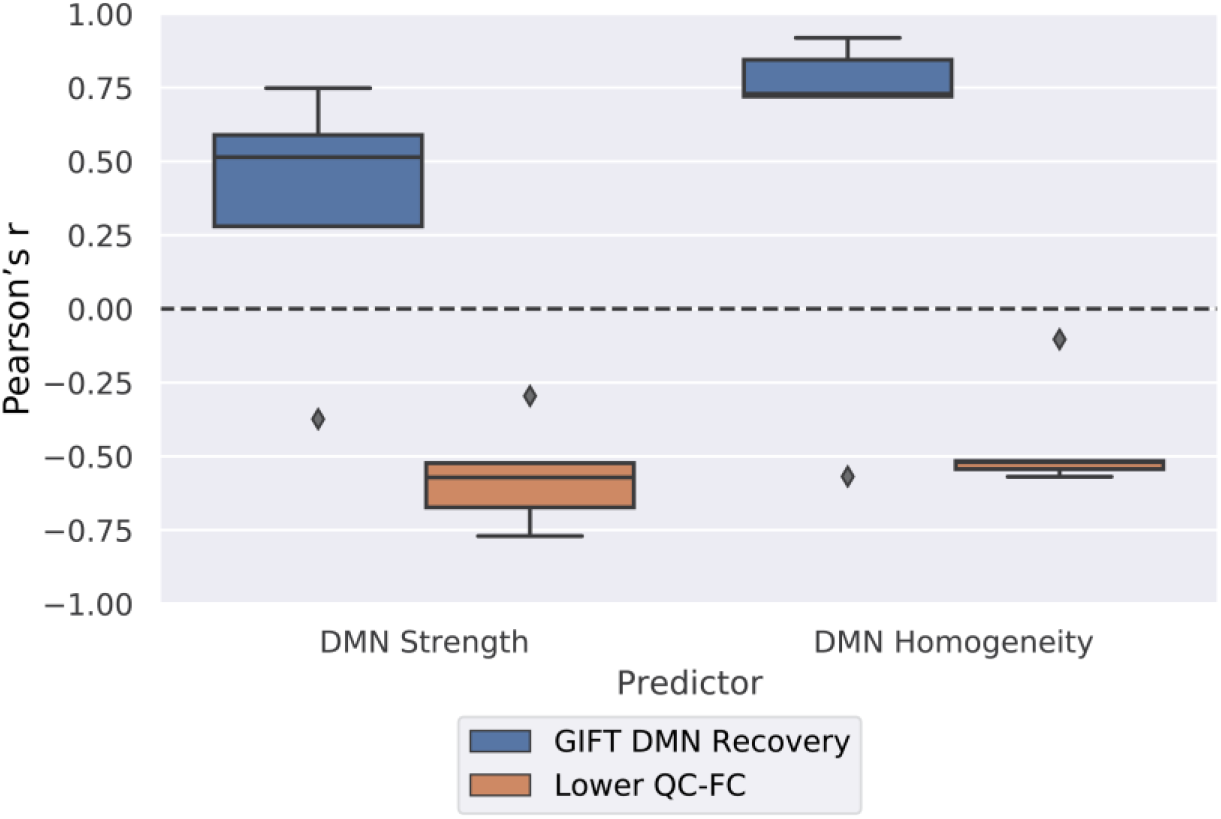
Investigation into the discrepancy between promising QC-FC yet poor RSN recovery for the fully concatenated [HMP, AROMA, Physio, Freq] pipeline. To probe the validity of the QC-FC metric, Pearson’s correlation was computed between measures of DMN quality and the QC-FC metric across the 18 pipelines. Shown here is the distribution of the resulting correlations for all disease datasets. A Positive correlation would indicate that higher DMN quality improves pipeline QC-FC, and vice versa. Correlations of DMN quality with GIFT (GICA) DMN recovery are also included for comparison. DMN Strength is measured by taking the mean of the FC values across all DMN edges (connections) within a subject, and then the grand mean across all subjects for a pipeline. DMN Homogeneity is measured by taking the negative standard-deviation of the FC values across all DMN edges within a subject, and then the mean of standard-deviations across all subjects for a pipeline. QC-FC values were negated to indicate improvement with increasing values. Results show that increases in DMN quality improves GICA Dice similarity scores while worsening QC-FC, indicating that QC-FC scores improve with the removal of DMN signal and bringing the metric’s validity into question.

### Closer examination of the modularity quality metric

While the Louvain algorithm automatically finds the FC network communities that optimize modularity quality, we sought to identify *what* networks were actually found and how they related to the canonical RSNs. We hypothesized that the communities which optimize modularity quality may not necessarily be biologically relevant or related to the canonical RSNs. They may instead be affected by motion artifact, as suggested by the [HMP, AROMA, Physio, Freq] pipeline having worse modularity quality-motion correlations than baseline. **Figure 9a** compares the mean number of communities found by the Louvain algorithm in the images processed with each pipeline. The [HMP, AROMA, Physio, Freq] pipeline (highlighted in the purple box, **Figure 9a**), which had the highest modularity quality *Q* in **Figure 7c**, also had the fewest number of communities. The Louvain algorithm found a mean of only 2.8 communities for this pipeline, while over 3 communities were found from *every* other pipeline. The [HMP, AROMA, Physio, Freq] pipeline also manifests as an outlier when the Dice similarity was computed between the canonical DMN and the most similar community from the Louvain result of each image (**Figure 9b**,**c**). The communities identified in the [HMP, AROMA, Physio, Freq] pipeline had the lowest Dice similarity with the DMN, indicating that data from this pipeline was not able to recover and distinguish well the primary RSN.

**Figure 9.**
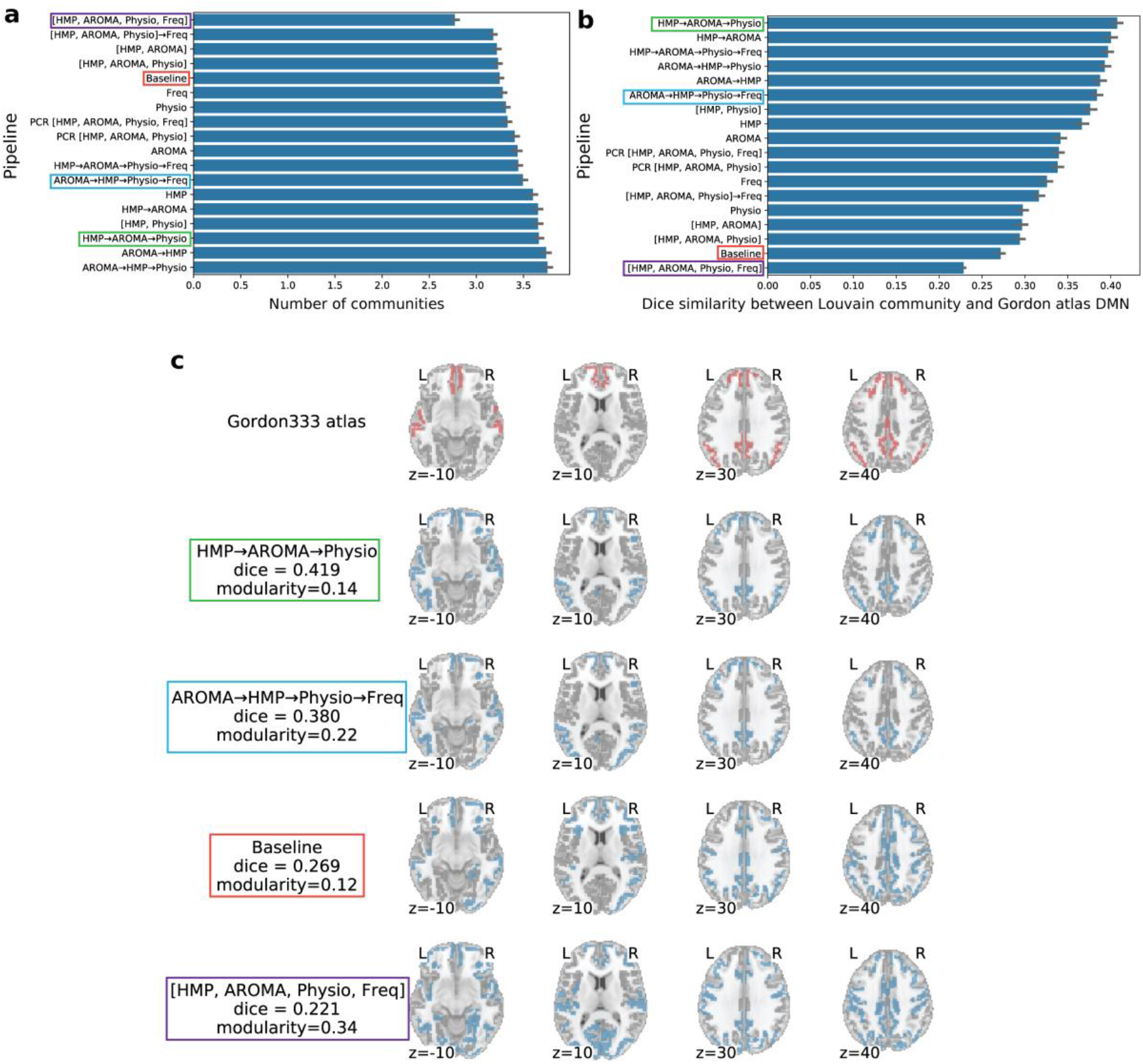
Investigation into the discrepancy between promising modularity yet poor RSN recovery for the fully concatenated [HMP, AROMA, Physio, Freq] pipeline. For this analysis, the number of communities computed by the Louvain algorithm for all 818 subjects in the ABIDE, EMBARC, and PPMI datasets was examined (a). We found this pipeline had the lowest number of communities, indicating less separability of the FC into distinct networks compared to other pipelines. DMN signal recovered by the Louvain algorithm was also examined using a similar template-matching approach as used in GICA, using the DMN template from the Gordon333 atlas. Dice score between the best matching Louvain community and the atlas for all subjects are shown in (b), where notably the [HMP, AROMA, Physio, Freq] achieves the lowest dice, indicating that the Louvain algorithm is unable to recover a DMN. A representative subject from the EMBARC dataset is shown in (c) to compare the Louvain partitions on different pipeline outputs with the Gordon333 atlas. DMN regions in the Gordon333 atlas are colored in red. For each pipeline, the Louvain partition most similar to the DMN is colored in blue. A comparatively high modularity *Q* for [HMP, AROMA, Physio, Freq] suggests that the modularity metric can give spurious high values on outputs that remove putative reliable true networks such as the DMN.

## Discussion

### Pipeline recommendations

Ciric et al. suggest that the optimal motion correction strategy will differ by metric, and that largely holds true with these results (Ciric et al. 2017). However, by prioritizing metrics that appear more reliable, we can begin to make some general recommendations. With reproducibility being a key desideratum in fMRI research, we first consider the HCP test-retest reliability (TRT) results. Based on TRT metrics for comparing the same subject’s FC, ReHo, fALFF, and individual DMN spatial maps under conditions of high and low motion, HMP→AROMA→Physio→Freq performed the best overall. In the GICA-based, group-level DMN maps, this fully sequential pipeline also recovers reproducible DMN maps between low- and high-motion images. Also, in the disease datasets, this sequential pipeline showed reasonable performance in recovering the DMN via GICA and SBC, ranking neither best nor worst. Overall, the HMP→AROMA→Physio→Freq shows the most promise for improving TRT of connectivity and RSN analyses in images affected by motion in varying degrees, and it is a recommended choice for removing motion artifacts in images of subjects with neurological and psychiatric disorders.

Adopting the geometrical perspective on nuisance regression suggested by Lindquist et al. 2019 may shed light on why the HMP→AROMA→Physio→Freq worked so well across several metrics and datasets. It may be that the reintroduction of artifact removed in previous steps of the sequential regression is *outweighed* by the efficacy of relatively small design matrices to fit the noise components in the data at each step. It could also be that previous regression steps may remove some actual neuronal signal that is reintroduced in later steps. Thus, although theoretically superior, concatenated regression may in many cases be inferior to sequential regression in practice. These speculations suggest a promising line of inquiry that future experiments could address.

### Concatenated regression

Lindquist et al. postulated that a single nuisance regression step, with all regressors concatenated in the same design matrix, would outperform sequential regression steps by avoiding reintroduction of artifacts (Lindquist et al. 2019). However, our results showed that the fully concatenated [HMP, AROMA, Physio, Freq] pipeline performed the poorest across the TRT metrics and DMN metrics, often showing even worse metrics than the uncorrected baseline images. Qualitative comparisons of DMN spatial maps indicated an apparent loss of RSN signal and the presence of remaining noise. These problems likely stem from fitting a GLM on a large design matrix containing 100s of regressors. *HMP* provides 24 regressors (after expansion with squares and derivatives), *AROMA* may create as many as 50 regressors (depending on the number of ICA components automatically selected by the AROMA algorithm), *Physio* contributes 8 regressors (with squares and derivatives), and *Freq* may contain as many as 135 regressors (depending on the number of sinusoids needed for the specified bandpass, which is determined by the granularity of frequencies needed to approximate the bandpass and by the number of volumes). The high dimensionality of the concatenated design matrix leads to two obstacles: 1) a great loss of degrees of freedom in the cleaned data and 2) an ill-posed GLM. The image timeseries analyzed in this work contained from 128 to 300 volumes, meaning that in the worst case, there would be more regressors than observations when fitting the GLM. Indeed, poor GLM fitting on the concatenated design matrix is shown by computing the partial coefficient of determination for each regressor type using the method described by (Bianciardi et al. 2009) (**Figure S14**). We conclude that the theoretical benefits of single-step concatenated regression, as proposed by Lindquist et al., do not appear to outweigh the practical obstacles of performing such a regression, at least at the current clinical TR of 1-2 seconds.

However, we still find evidence that future work aimed at optimizing such a single-step, concatenated regression may eventually prove fruitful. For example, applying PCR to reduce the dimensionality of the concatenated design matrix, before performing the regression was substantially more effective than using the full design matrix. The PCR approach substantially improved performance in the FC and DMN TRT metrics on the HCP dataset and the SBC DMN Dice similarity on the disease datasets. Increasing the degree of dimensionality reduction improved performance further, though not to the extent of the sequential pipelines (**Figure S15**). Another alternative is the [HMP, AROMA, Physio] →Freq pipeline, with Freq regression performed separately since it contains the most regressors. This approach performed nearly as well as the leading HMP→AROMA→Physio→Freq pipeline in the DMN Dice similarity metrics on the disease datasets.

### Validity of motion correction metrics

Interestingly, despite performing poorly in TRT and DMN metrics, the fully concatenated pipeline showed the best QC-FC correlation and modularity quality. However, further analysis demonstrated serious shortcomings of the QC-FC correlation and modularity quality metrics. Lower (better) QC-FC correlation was associated with lower DMN FC strength, indicating that QC-FC correlation is biased towards pipelines that *suppressed* RSN activity. Similarly, lower QC-FC correlation was associated with lower DMN homogeneity, i.e. higher variance in DMN FC. This suggests a case of regression attenuation, where higher variance in the data biases Pearson’s correlation towards zero (Saccenti et al. 2020; Thouless 1939). In other words, a pipeline could hypothetically achieve a good QC-FC correlation by simply reducing FC to high-variance noise. Further evidence of this signal loss when measuring the temporal signal-to-noise ratios of these pipelines (**Figure S16**). Since the QC-FC-dd metric is derived from the QC-FC metric, its reliability as an index of motion artifact removal is doubtful as well. Therefore, prioritizing QC-FC and QC-FC-dd correlations as motion correction metrics may inadvertently lead to removing true biological signals from the data, and we do not recommend using QC-FC and QC-FC-dd correlations in their current form.

The modularity quality metric, *Q*, also raised concerns. Ciric et al. showed that modularity quality can be negatively correlated with subject motion; for some motion correction approaches, this correlation was as large as *r* = −0.494 (Ciric et al. 2017). This is undesirable, because ideally a subject’s network modularity should be decoupled from and unassociated with their motion level. An additional concern comes from the use of the Louvain algorithm to compute modularity quality. This algorithm performs a greedy optimization to identify graph communities that maximize modularity quality, but there is no guarantee that these communities are meaningful or related to actual RSNs. Our results showed that the [HMP, AROMA, Physio, Freq] pipeline with the best modularity quality also had correlation between modularity quality and motion that was worse than baseline and significantly fewer communities than the other pipelines. The Louvain-partitioned communities did not appear to reflect any known RSNs, such as the highly reproducible DMN. Also, there is no clear neurophysiological basis for the optimal value of modularity quality. The “true” modularity of the human brain is unknown, meaning that a higher modularity is not clearly better. When developing motion correction methods, we recommend that researchers prioritize metrics that have a strong neurophysiological basis, such as the identifiability of canonical RSNs, over blind partitioning methods like the Louvain algorithm.

Most importantly, we advocate that motion correction methods be evaluated with a variety of metrics and on several datasets. While many previous studies have focused on metrics based on FC (Ciric et al. 2017; Parkes et al. 2018; Power et al. 2015), FC is only a single method of rs-fMRI analysis. To fully understand the efficacy of a given motion correction method, one should also consider RSN-based measures (e.g. GICA, SBC, spatial map integrity) and local connectivity (e.g. ReHo, fALFF), in the context of disease as well as test-retest reliability.

### Limitations

This is work is not intended to be an exhaustive comparison of motion correction approaches. We considered only a subset of the many nuisance regressor types proposed by the community and a selection of the possible regressor sequences, as our main goal was the comparison of concatenated with sequential regression approaches. In future work, comparisons of additional nuisance regressor generation methods such as ANATICOR or ICA-FIX could be made (Jo et al. 2013; Salimi-Khorshidi et al. 2014). In the meantime, our analysis included some of the most common nuisance regressors used in the fMRI research community.

We have not considered global signal regression (GSR). Though GSR has been suggested to reduce QC-FC correlation (Ciric et al. 2017), there is much uncertainty among fMRI researchers about whether GSR may introduce artifactual anticorrelations or remove biological signal (Murphy et al. 2009; Liu et al. 2017). Also, we have shown that conclusions based on the QC-FC correlation metric may be unreliable. Testing of GSR with additional metrics, such as TRT and DMN recovery, may be more informative.

Finally, we did not examine despiking or scrubbing (volume censoring) methods. These methods, which are much more aggressive than the HMP, AROMA, and Physio regression steps tested here, involve the truncation or removal of data from the fMRI timeseries. While despiking and scrubbing can be effective (Jo et al. 2013; Power et al. 2012; Satterthwaite et al. 2013), they may impede group-level analyses such as GICA unless the censored volumes are appropriately interpolated. Consequently, we leave these comparisons for future work.

## Conclusion

This work quantitatively compares motion correction strategies, consisting of extensive combinations of regressors in sequence or in concatenation and provides new insights and recommendations for motion correction. Through this analysis of multiple metrics across an array of healthy and diseased datasets, the work makes the following contributions. *First*, critical limitations in multiple commonly used motion correction metrics, including modularity quality and QC-FC and QC-FC-dd correlations, are identified. It was demonstrated that these metrics are prone to mislead researchers to select motion correction approaches that do not recover RSN signals or that even remove true biological signals from fMRI. As an alternative, a set of metrics based on neurophysiological priors and reproducibility is recommended. *Second*, limitations in a fully concatenated regression approach are demonstrated and are recommended to be addressed in further development in high-dimensionality regression approaches. *Third*, a pipeline, the sequential HMP → AROMA →Physio → Freq pipeline is recommended as it is shown to be robust and outperform other sequential and concatenation pipelines across multiple datasets. This work is intended to help other fMRI researchers to select appropriate motion correction strategies, to evaluate such strategies more critically, and to further the analysis of fMRI data that reflects true biological signal.

## Acknowledgements

We thank Cooper Mellema and Alex Treacher for their feedback during the writing of this manuscript.

Data were provided in part by the Human Connectome Project, WU-Minn Consortium (Principal Investigators: David Van Essen and Kamil Ugurbil; 1U54MH091657) funded by the 16 NIH Institutes and Centers that support the NIH Blueprint for Neuroscience Research; and by the McDonnell Center for Systems Neuroscience at Washington University.

PPMI-–a public-private partnership-–is funded by the Michael J. Fox Foundation for Parkinson’s Research and funding partners, including Abbvie, Allergan, Avid Radiopharmaceuticals, Biogen, Biolegend, Bristol-Myers Squibb, Celgene, Denali, GE Healthcare, Genentech, GlaxoSmithKline, Lilly, Lundbeck, Merck, Meso Scale Discovery, Pfizer, Piramal, Prevail Therapeutics, Roche, Sanofi Genzyme, Servier, Takeda, Teva, UCB, Verily, and Voyager Therapeutics.

## Data availability statement

This work uses publicly available functional MRI data from the Human Connectome Project (Young Adult S1200 release, https://www.humanconnectome.org/study/hcp-young-adult), the Parkinson Progression Markers Initiative (www.ppmi-info.org/data), the Autism Brain Imaging Data Exchange (http://fcon_1000.projects.nitrc.org/indi/abide/), and the Establishing Moderators and Biosignatures of Antidepressant Response for Clinical Care study (http://embarc.utsouthwestern.edu/).

## Footnotes

1 www.ppmi-info.org/data

2 https://brainnexus.com/resting-state-fmri-templates/

## Notes

### Competing Interest Statement

The authors have declared no competing interest.

