## Supplemental Material for "Pitfalls and recommended strategies and metrics for suppressing motion artifacts in functional MRI"

**Table S1**. Resting-state fMRI acquisition parameters for each dataset. From ABIDE, analysis included data from the New York University (NYU), Georgetown University (GU), and Kennedy Krieger Institute (KKI) sites.

| | HCP | ABIDE | EMBARC | PPMI |
|---|---|---|---|---|
| **Field strength** | 3 T | 3 T | 3 T | 3 T |
| **Scanner(s)** | Siemens Connectome | Siemens Allegra (NYU), Siemens TrioTim (GU) Philips Achieva (KKI) | GE Signa HDx, Siemens TrioTim, Philips Achieva, Philips Ingenia | Siemens TrioTim |
| **Pulse sequence** | GE-EPI | GE-EPI | GE-EPI | GE-EPI |
| **TR** | 720 ms (downsampled to 2880 ms) | 2000 ms (NYU) 2000 ms (GU)<br>2500 ms (KKI) | 2000 ms | 2400 ms |
| **TE** | 33.1 ms | 15 ms (NYU)<br>30 ms (GU)<br>30 ms (KKI) | 28 ms | 25 ms |
| **Flip angle** | 52° | 90° (NYU)<br>90° (GU)<br>75° (KKI) | 90° | 80° |
| **Voxel size** | 2 mm isotropic | 3 x 3 x 4 mm (NYU)<br>3 x 3 x 3 mm (GU)<br>3 x 3 x 3 mm (KKI) | 3.2 x 3.2 x 3.1 mm | 3.3 mm isotropic |
| **Image voxel dimensions** | 104 x 90 x 72 | 64 x 80 x 33 (NYU)<br>64 x 64 x 43 (GU)<br>96 x 96 x 47 (KKI) | 64 x 64 x 39 | 68 x 66 x 33 |
| **Volumes** | 1200 (downsampled to 300) | 180 (NYU)<br>154 (GU)<br>128 (KKI) | 180 | 210 |
| **Total acquisition time** | 864 s | 360 s (NYU)<br>308 s (GU)<br>320 s (KKI) | 360 s | 504 s |

**Table S2**. Dataset demographics.

| | HCP | ABIDE | EMBARC | PPMI |
|---|---|---|---|---|
| **Subjects** | 50 | 359 | 309 | 150 |
| **Mean age (± s.d.)** | 28.9 ± 3.6 | 12.0 ± 2.5 | 37.0 ± 13.2 | 62.8 ± 10.1 |
| **Male** | 20 | 255 | 104 | 106 |
| **Female** | 30 | 104 | 205 | 44 |

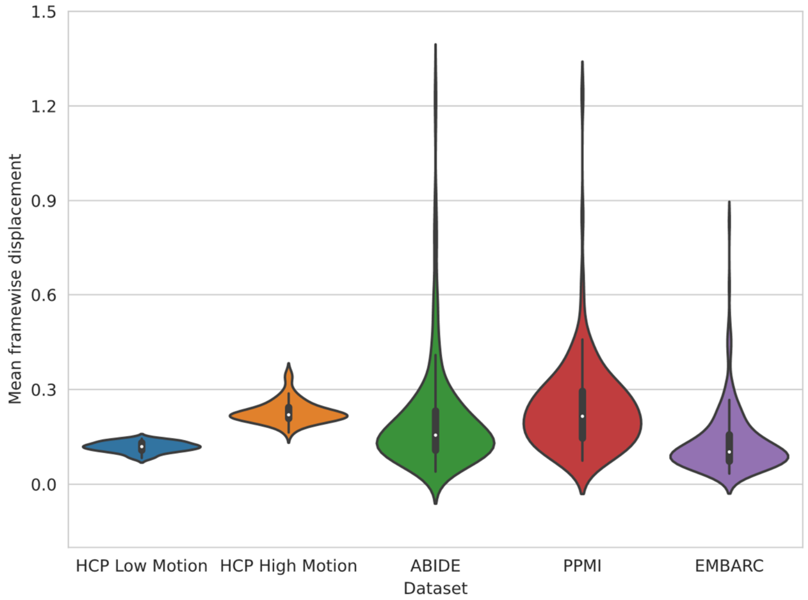


**Figure S1.** Mean framewise displacement distribution of each dataset.

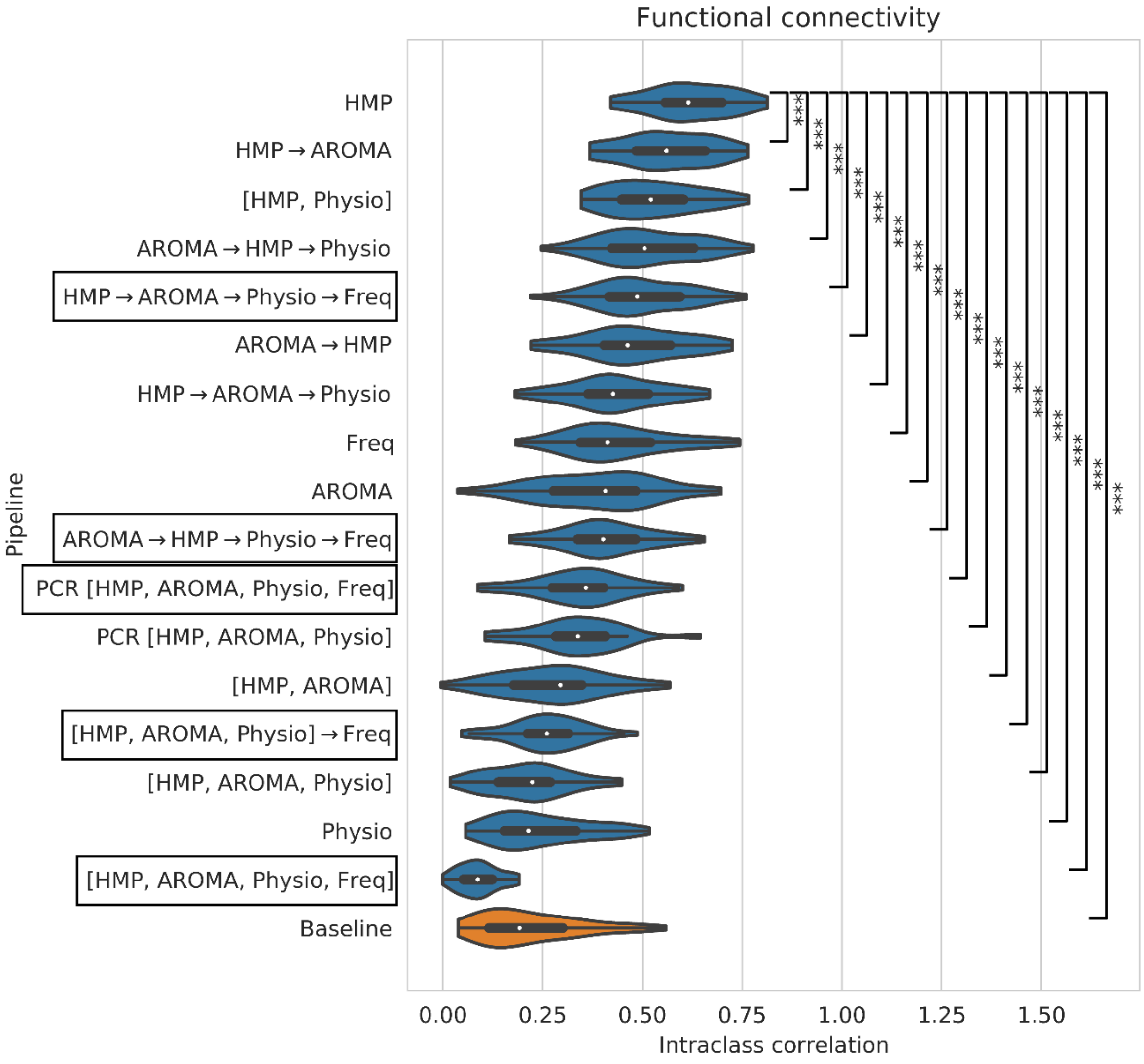


**Figure S2**. Test-retest reliability of *functional connectivity* computed from high- and low-motion HCP images acquired on the same day, after each motion correction pipeline. Functional connectivity was computed using the 333-region Gordon atlas. Significant differences compared to the top-ranking pipeline are indicated: * p < 0.05, ** p < 0.01, *** p < 0.001. Boxed pipelines, which contain all 4 regressor types, were compared in the main text results (Figure 3a).

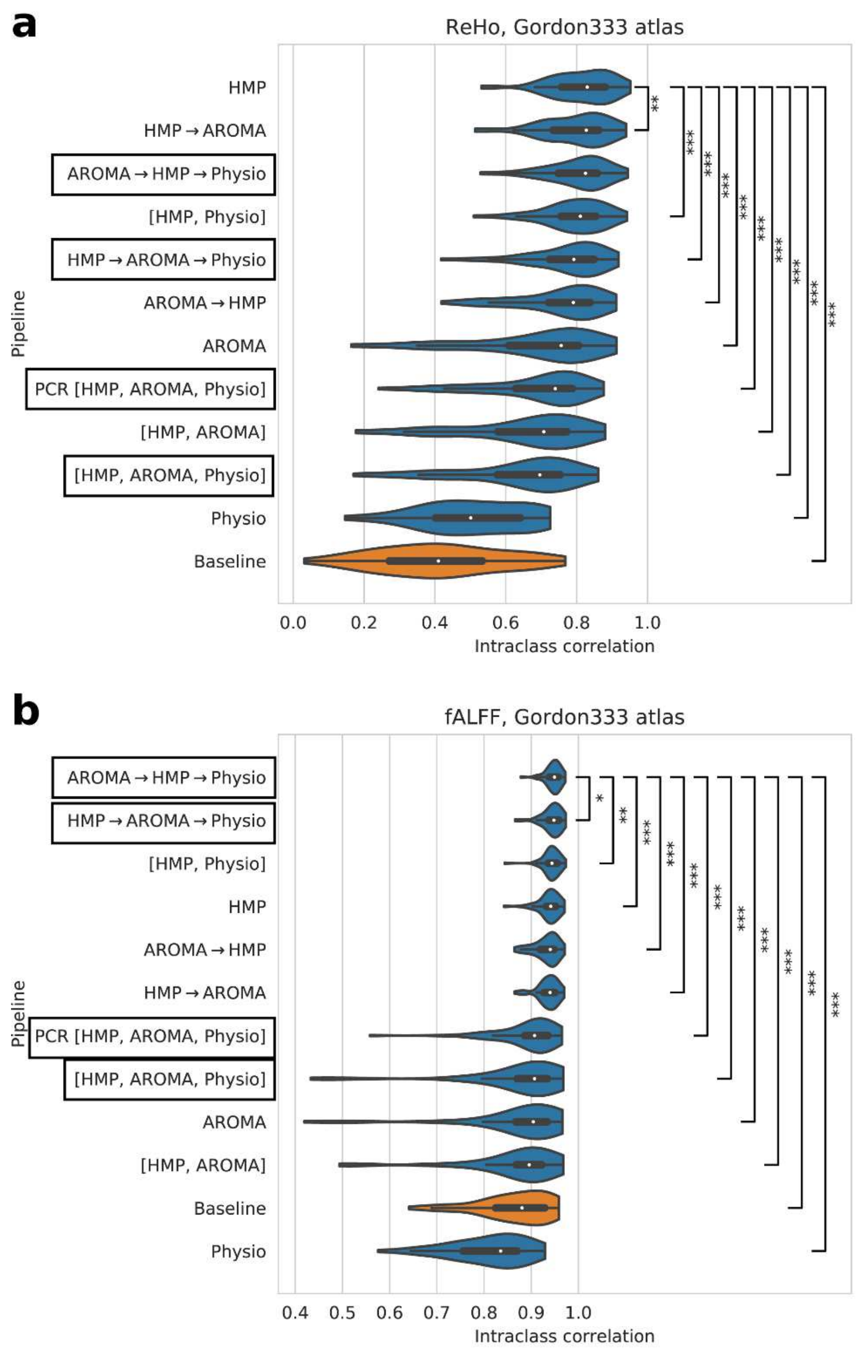


**Figure S3**. Test-retest reliability of *regional homogeneity (ReHo)* and *fractional amplitude of low frequency fluctuations (fALFF)* computed from high- and low-motion HCP images acquired on the same day, after each motion correction pipeline. Because the computation of ReHo and fALFF include predefined bandpass filters, pipelines that include Freq regressors were not included in this comparison. Significant differences compared to the top-ranking pipeline are indicated: * $p < 0.05$, ** $p < 0.01$, *** $p < 0.001$. Boxed pipelines, which contain all 4 regressor types, were compared in the main text results (Figure 3b-c).

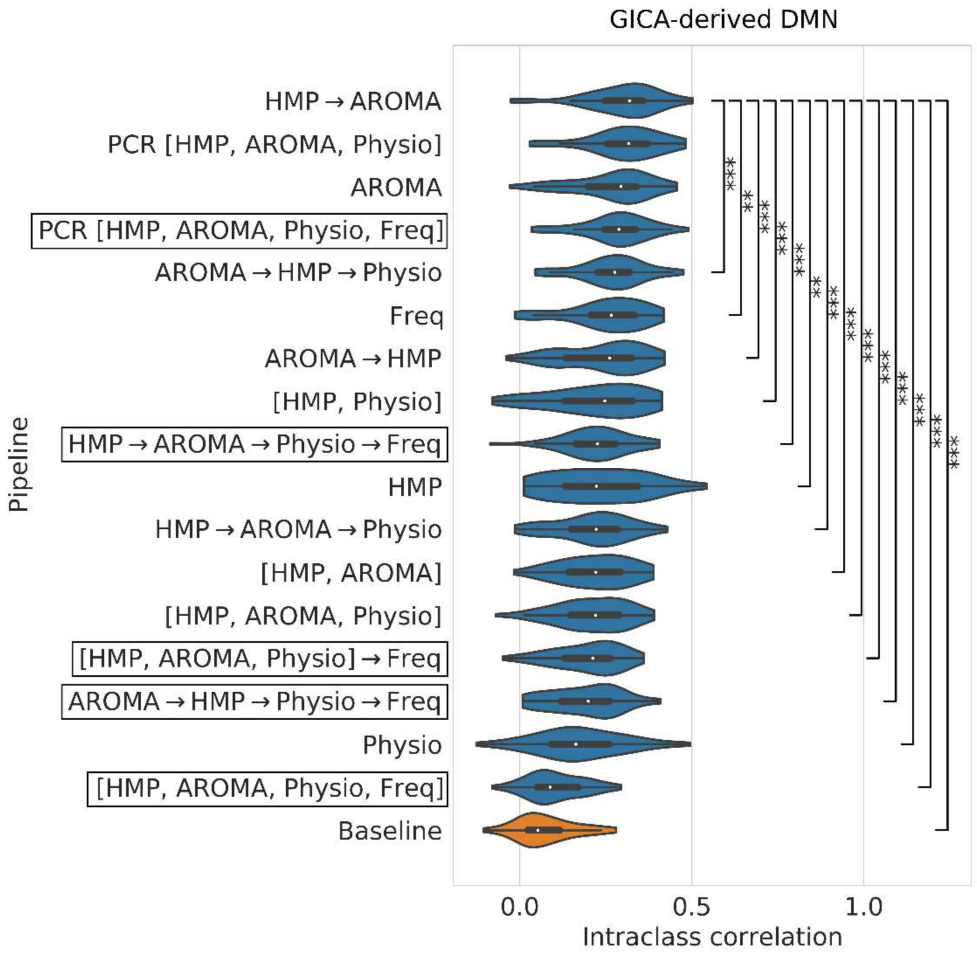


**Figure S4**. Test-retest reliability of default mode network maps computed using *group independent components analysis (GICA)* from high- and low-motion HCP images acquired on the same day, after each motion correction pipeline. Significant differences compared to the top-ranking pipeline are indicated: * $p < 0.05$, ** $p < 0.01$, *** $p < 0.001$. Boxed pipelines, which contain all 4 regressor types, were compared in the main text results (Figure 3d).

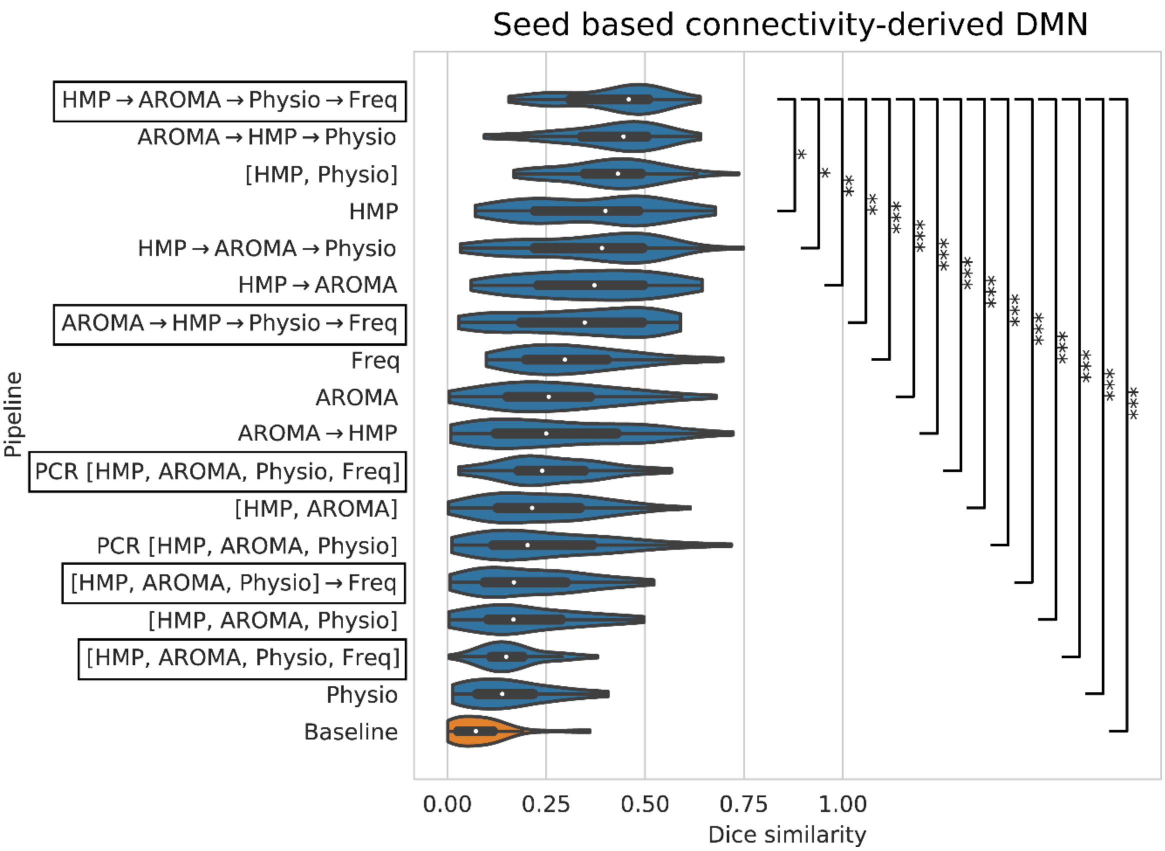


**Figure S5**. Test-retest reliability of default mode network maps computed using *seed-based connectivity* from high- and low-motion HCP images acquired on the same day, after each motion correction pipeline. Seed-based connectivity maps were generated using a 6 mm radius seed at the right posterior cingulate cortex at MNI coordinate (4, -54, 26). Significant differences compared to the top-ranking pipeline are indicated: * $p < 0.05$, ** $p < 0.01$, *** $p < 0.001$. Boxed pipelines, which contain all 4 regressor types, were compared in the main text results (Figure 3e).

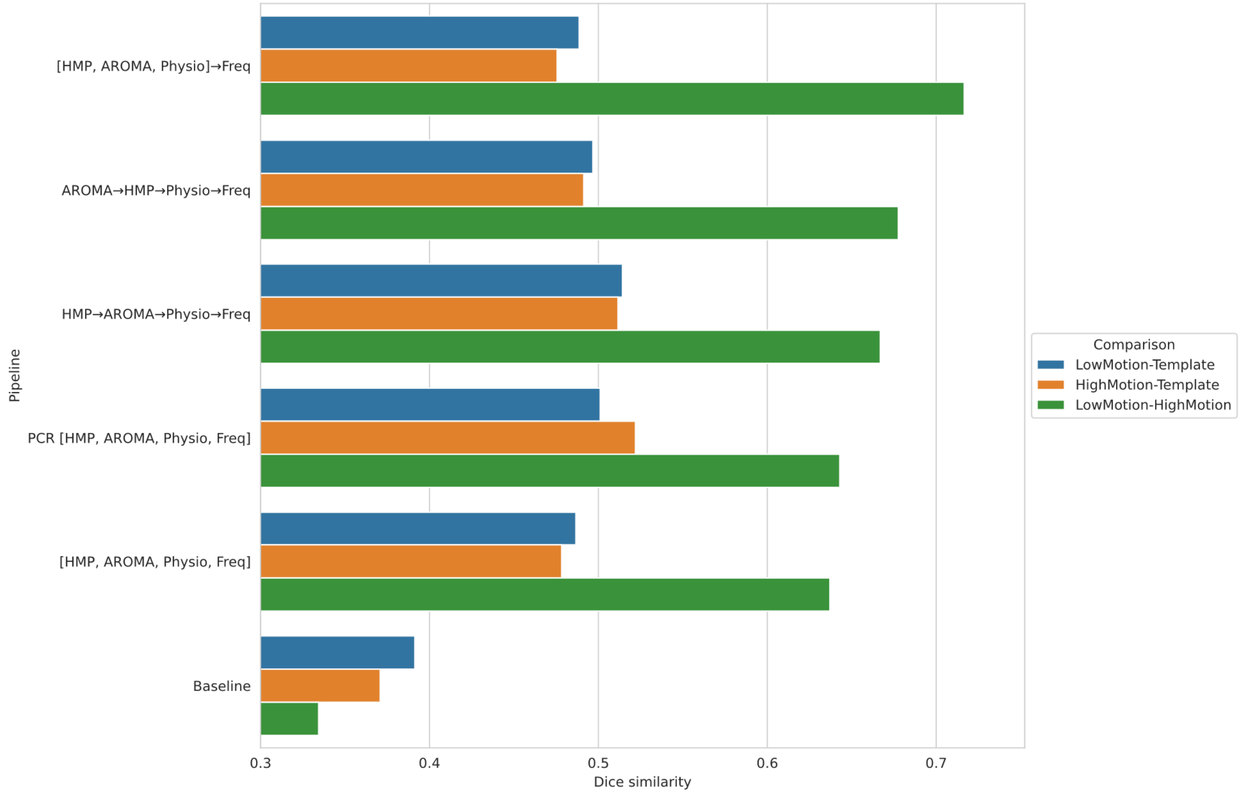


**Figure S6.** Dice similarity of group-level default mode network maps computed using *group independent components analysis (GICA)* from HCP images. Dice was computed between high-motion group map and the low-motion group map, between the high-motion group map and the Thomason et al. 2011 template, and between the low-motion group map and the template.

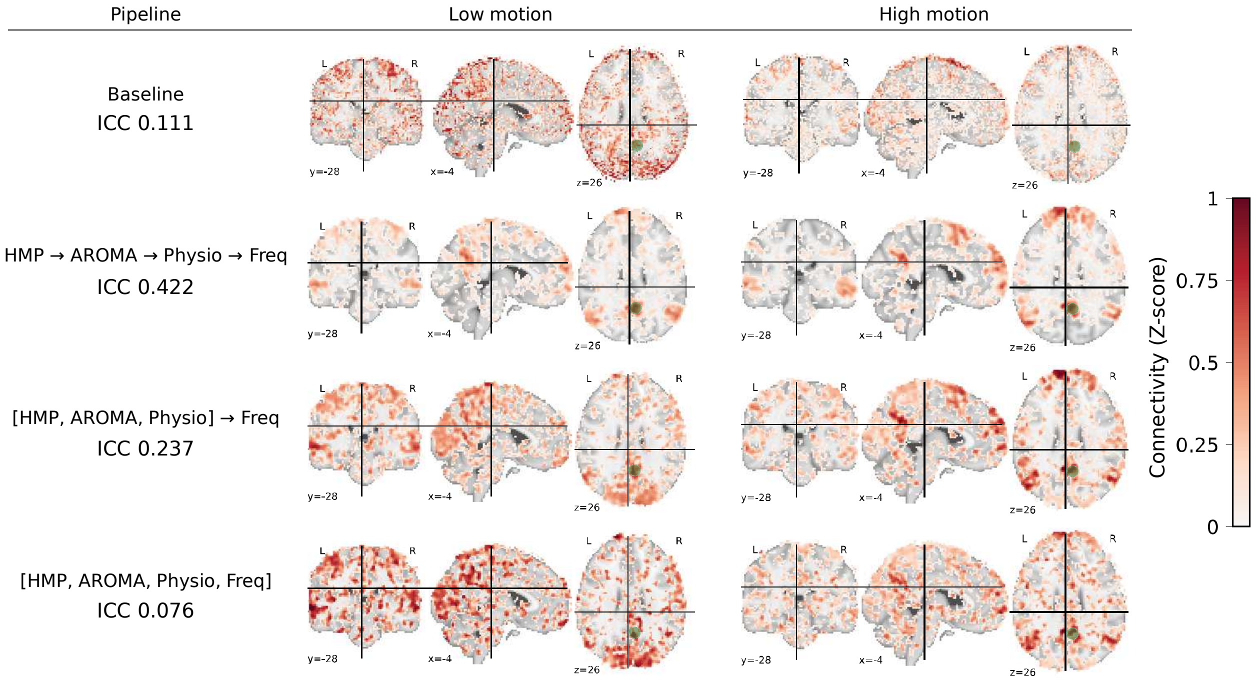


**Figure S7.** Representative example of an HCP subject's seed-based connectivity (SBC) maps computed from high- and low-motion images acquired the same day, after different motion correction pipelines. Test-retest reliability between the high- and low-motion SBC maps is compared across pipelines in main text Figure 3. SBC was computed using a 6 mm radius spherical seed in the right posterior cingulate cortex, shown in green, to recover the default mode network (DMN). Connectivity was computed using Pearson's correlation then Fisher transformed into Z-scores. For visualization purposes, SBC maps were thresholded at Z > 0 below. The un-corrected baseline SBC maps show substantial differences, while the HMP → AROMA → Physio → Freq and [HMP, AROMA, Physio] → Freq pipelines show better similarity between high- and low-motion maps. These SBC maps also better visualize the DMN, including the characteristic parietal, prefrontal, and hippocampal components.

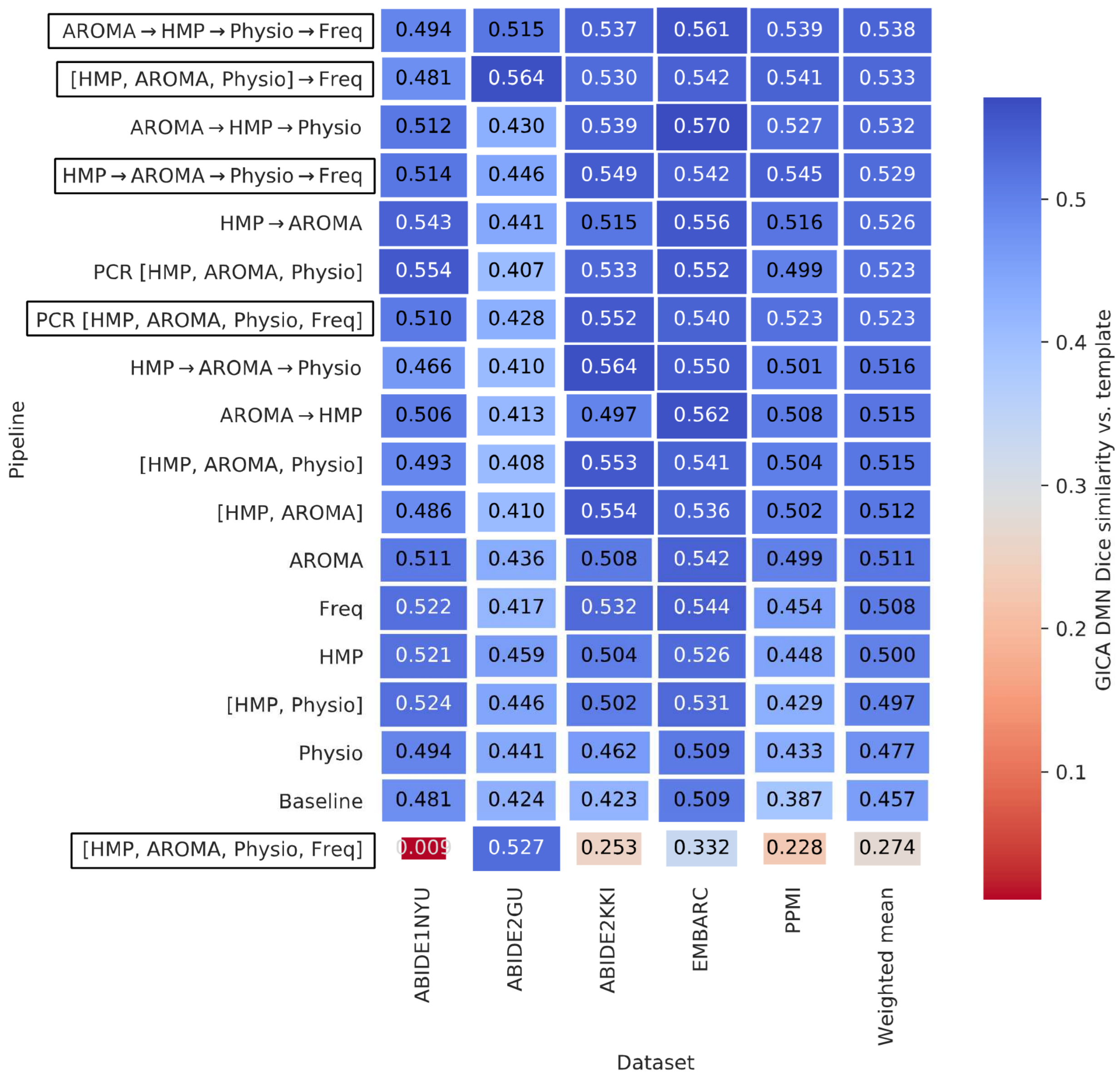


**Figure S8.** Values of *Dice similarity coefficient* between the brainnexus DMN template and the optimized *group ICA (GICA)* spatial map across high motion subjects for each pipeline (along the vertical axis) and dataset (along the horizontal axis). Darker blue colors and larger boxes indicate higher Dice score and better pipeline performance in recovering a DMN that matches the template. Pipelines are sorted by the sample size-weighted mean across all datasets (last column), with AROMA → HMP → Physio → Freq having the greatest mean. Boxed pipelines, which contain all 4 regressor types, were compared in the main text results (Figure 5a).

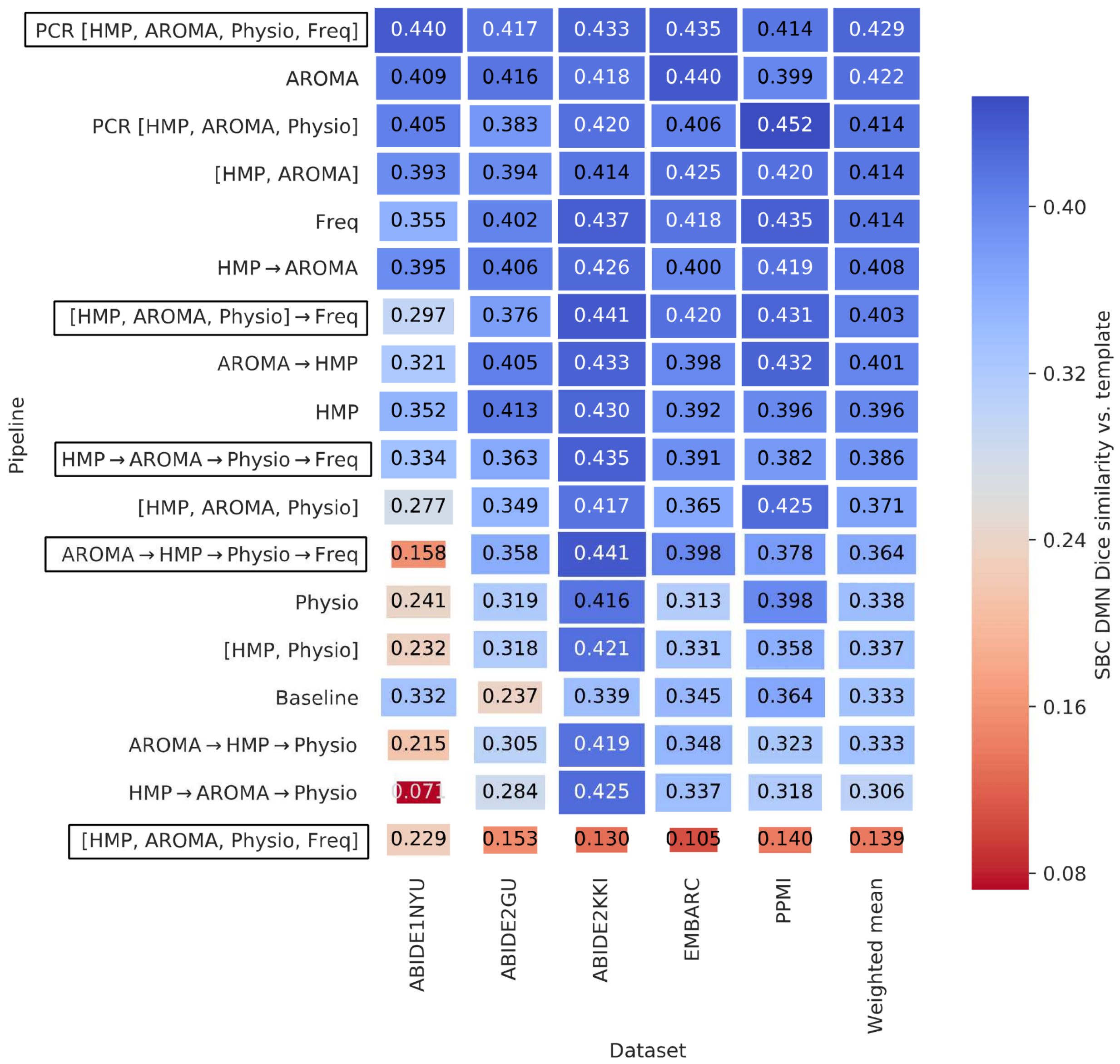


**Figure S9.** Values of *Dice similarity coefficient* between the Thomas et al. 2011 DMN template and the optimized *seed-based connectivity* map (seed at PCC) for each pipeline (along the vertical axis) and dataset (along the horizontal axis). Darker blue colors and larger boxes indicate higher Dice score and better pipeline performance in recovering a DMN that matches the template. Pipelines are sorted by the sample size-weighted mean across all datasets (last column), with PCR[HMP, AROMA, Physio, Freq] having the greatest mean. Boxed pipelines, which contain all 4 regressor types, were compared in the main text results (Figure 5b).

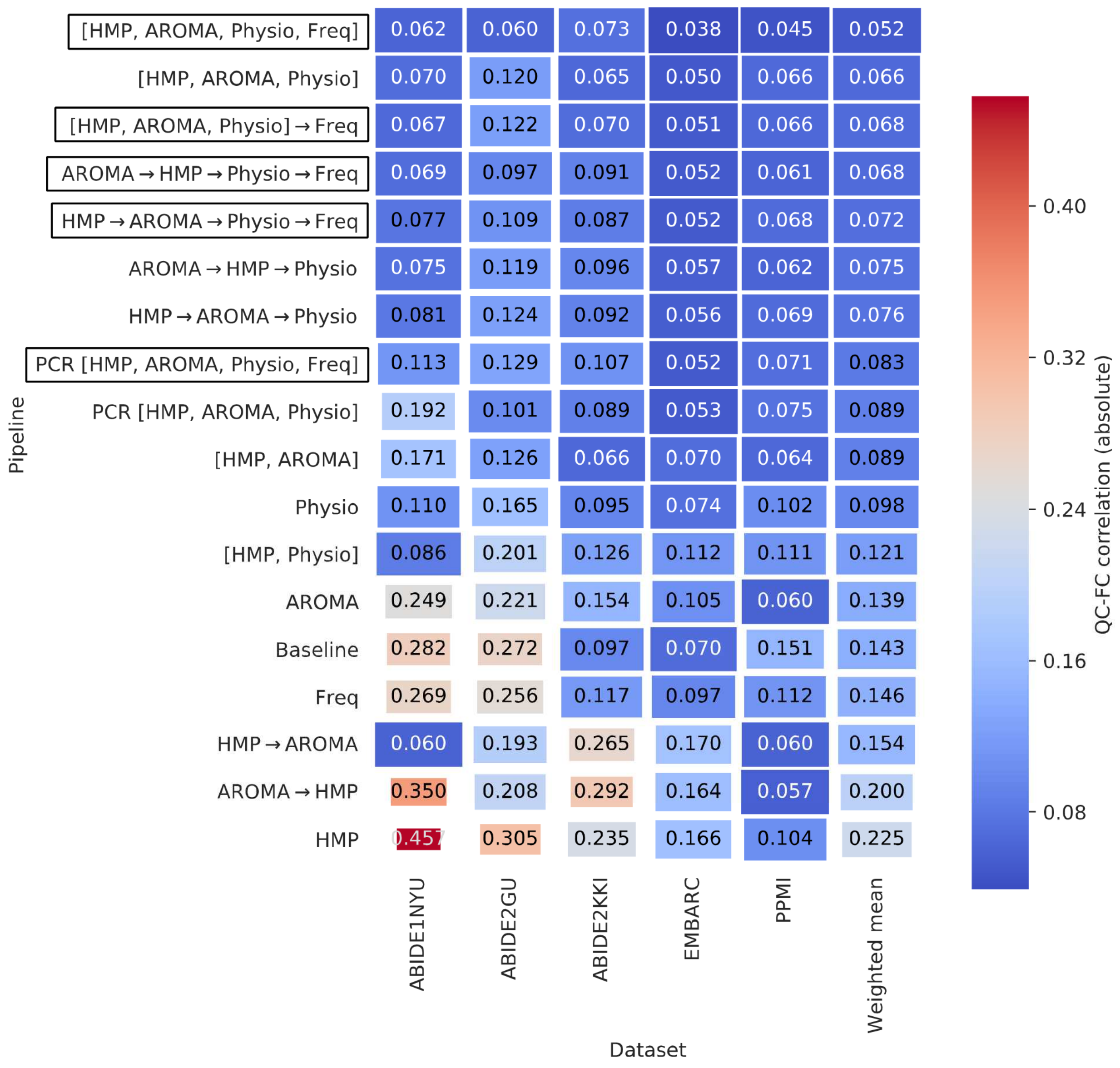


| Pipeline | ABIDE1NYU | ABIDE2GU | ABIDE2KKI | EMBARC | PPMI | Weighted mean |
|---|---|---|---|---|---|---|
| [HMP, AROMA, Physio, Freq] | 0.062 | 0.060 | 0.073 | 0.038 | 0.045 | 0.052 |
| [HMP, AROMA, Physio] | 0.070 | 0.120 | 0.065 | 0.050 | 0.066 | 0.066 |
| [HMP, AROMA, Physio] → Freq | 0.067 | 0.122 | 0.070 | 0.051 | 0.066 | 0.068 |
| AROMA → HMP → Physio → Freq | 0.069 | 0.097 | 0.091 | 0.052 | 0.061 | 0.068 |
| HMP → AROMA → Physio → Freq | 0.077 | 0.109 | 0.087 | 0.052 | 0.068 | 0.072 |
| AROMA → HMP → Physio | 0.075 | 0.119 | 0.096 | 0.057 | 0.062 | 0.075 |
| HMP → AROMA → Physio | 0.081 | 0.124 | 0.092 | 0.056 | 0.069 | 0.076 |
| PCR [HMP, AROMA, Physio, Freq] | 0.113 | 0.129 | 0.107 | 0.052 | 0.071 | 0.083 |
| PCR [HMP, AROMA, Physio] | 0.192 | 0.101 | 0.089 | 0.053 | 0.075 | 0.089 |
| [HMP, AROMA] | 0.171 | 0.126 | 0.066 | 0.070 | 0.064 | 0.089 |
| Physio | 0.110 | 0.165 | 0.095 | 0.074 | 0.102 | 0.098 |
| [HMP, Physio] | 0.086 | 0.201 | 0.126 | 0.112 | 0.111 | 0.121 |
| AROMA | 0.249 | 0.221 | 0.154 | 0.105 | 0.060 | 0.139 |
| Baseline | 0.282 | 0.272 | 0.097 | 0.070 | 0.151 | 0.143 |
| Freq | 0.269 | 0.256 | 0.117 | 0.097 | 0.112 | 0.146 |
| HMP → AROMA | 0.060 | 0.193 | 0.265 | 0.170 | 0.060 | 0.154 |
| AROMA → HMP | 0.350 | 0.208 | 0.292 | 0.164 | 0.057 | 0.200 |
| HMP | 0.457 | 0.305 | 0.235 | 0.166 | 0.104 | 0.225 |

**Figure S10.** Values of *median QC-FC correlations* of the 55,278 FC matrix edges represented by color for each pipeline (along the vertical axis) and dataset (along the horizontal axis). Darker blue colors and larger boxes indicate lower FC motion contamination and better pipeline performance, while lighter colors and smaller boxes indicate worse pipeline performance. Pipelines are sorted by sample size-weighted mean across all datasets (last column), with [HMP, AROMA, Physio, Freq] ranking as best pipeline followed by a mix of sequential and concatenated pipelines. Boxed pipelines, which contain all 4 regressor types, were compared in the main text results (Figure 7a).

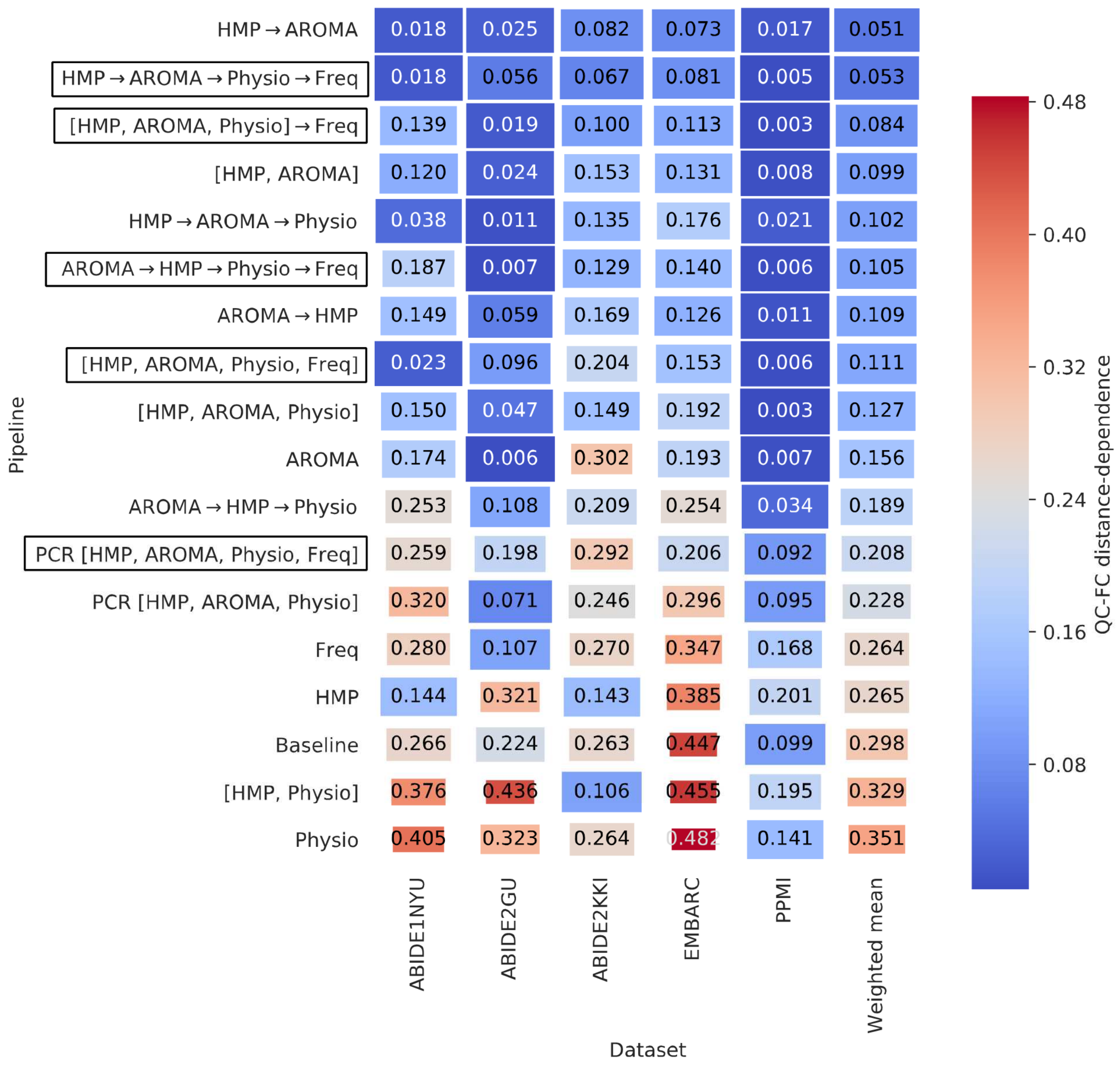


**Figure S11.** Values of *QC-FC distance dependence* for each pipeline (along the vertical axis) and dataset (along the horizontal axis). Darker blue colors and larger boxes indicate lower distance-dependent artifacts and better pipeline performance, while lighter colors and smaller boxes indicate worse pipeline performance. Pipelines are sorted by the sample size-weighted mean across all datasets (last column), with HMP→AROMA ranking as best pipeline, with HMP → AROMA → Physio → Freq ranking second, followed by a mix of sequential and concatenated pipelines. Boxed pipelines, which contain all 4 regressor types, were compared in the main text results (Figure 7b).

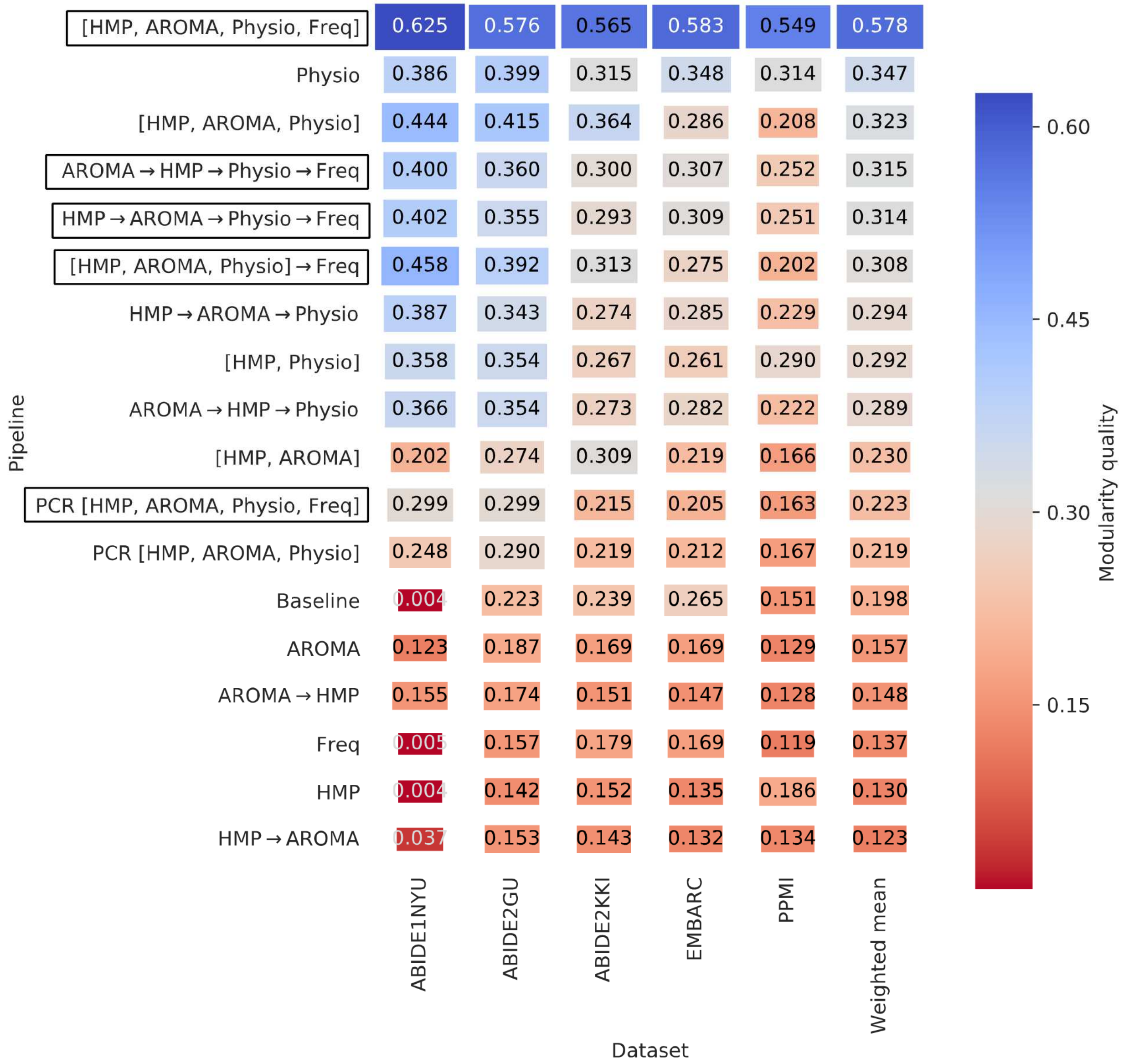


**Figure S12.** Values of *Modularity quality* for each pipeline (along the vertical axis) and dataset (along the horizontal axis). Darker blue colors and larger boxes indicate greater FC network modularity. Pipelines are sorted by the sample size-weighted mean across all datasets (last column), with [HMP, AROMA, Physio, Freq] having the highest mean. This metric is difficult to interpret, as network modularity may be enhanced by structured noise as well. Boxed pipelines, which contain all 4 regressor types, were compared in the main text results (Figure 7c).

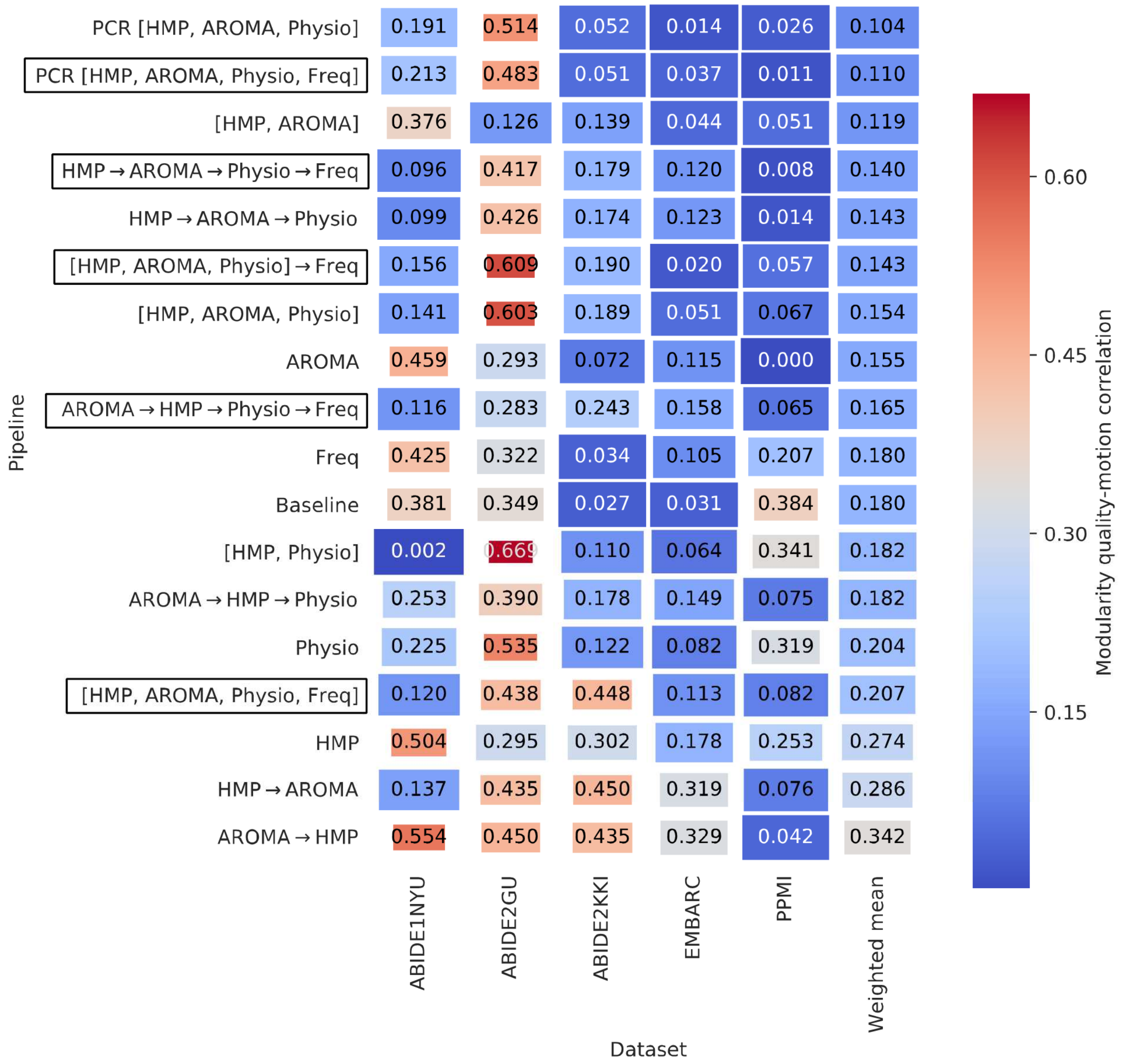


**Figure S13.** Values of *modularity quality-motion correlation* (Spearman's rank correlation) for each pipeline (along the vertical axis) and dataset (along the horizontal axis). Darker blue boxes and larger boxes indicate lesser motion contamination of the Modularity quality metric. Pipelines are sorted by the sample size-weighted mean across all datasets (last column), with PCR[HMP, AROMA, Physio] having the lowest contamination. Boxed pipelines, which contain all 4 regressor types, were compared in the main text results (Figure 7d).

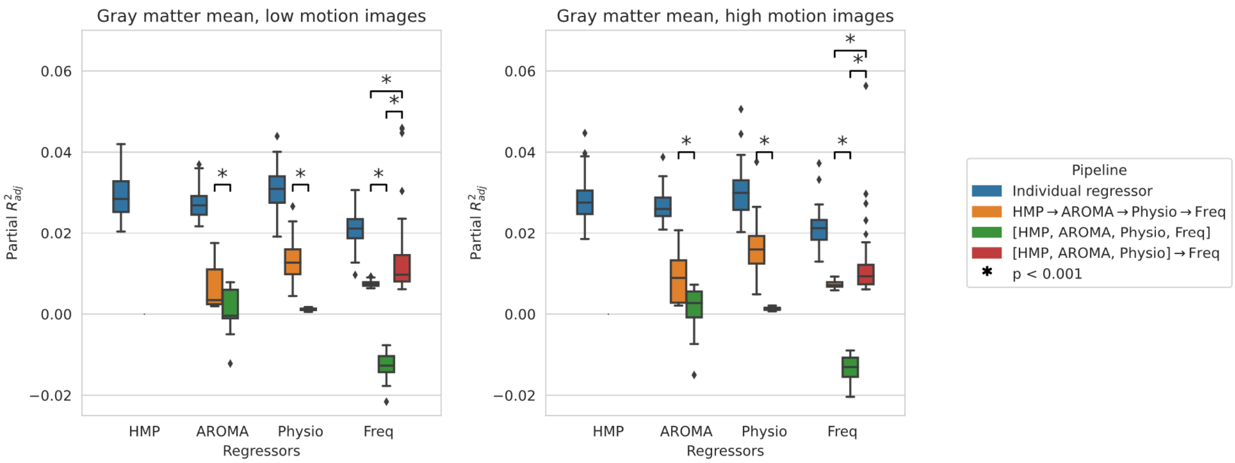


**Figure S14**. Adjusted partial $R^2$, measuring the fMRI signal variance explained by each regressor type in the HCP dataset. This metric was computed for each image using the approach of Bianciardi et al. 2009, then averaged over all gray matter voxels. Compared here are the regressors types applied individually to the raw images (blue), the sequential HMP → AROMA → Physio → Freq pipeline (orange), the concatenated [HMP, AROMA, Physio, Freq] pipeline (green), and the mixed pipeline with frequency regressors applied in a separate step (red). Note that the higher partial $R^2$ of the regressors applied individually compared to in combination indicate that the regressor types are somewhat collinear.

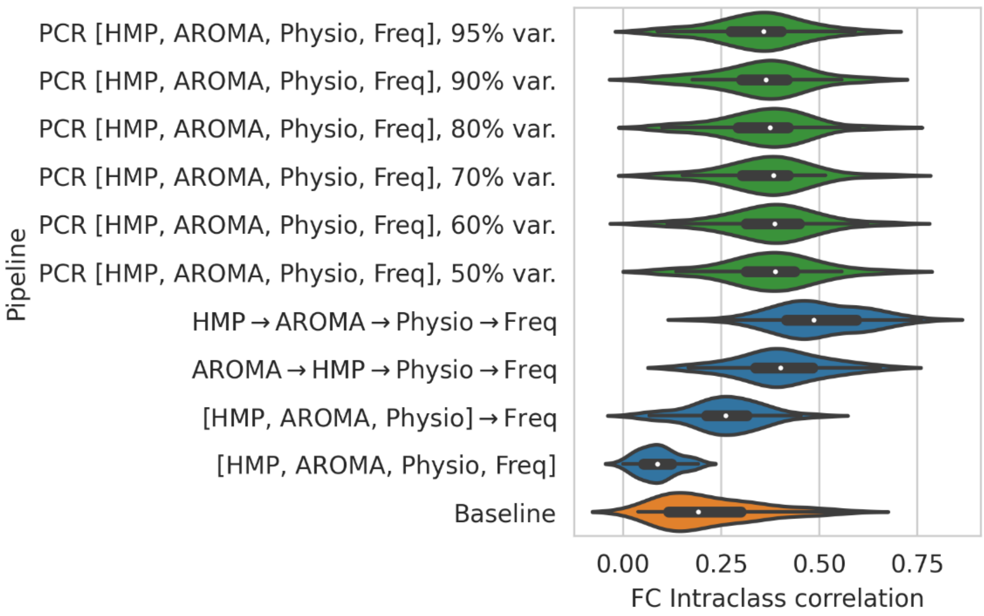


**Figure S15.** Comparison of principal components regression (PCR) of the concatenated design matrix while varying the number of retained principal components. The degree of dimensionality reduction was varied from 95% variance explained (more components retained) to 50% variance explained (fewer components retained). After applying each approach to the HCP dataset, test-retest reliability of functional connectivity was computed. Pipelines compared in this figure include the PCR variations (green), the other pipelines in the main results (blue), and the uncorrected baseline image (orange). The main results (Figure 3) include the PCR with 95% variance explained.

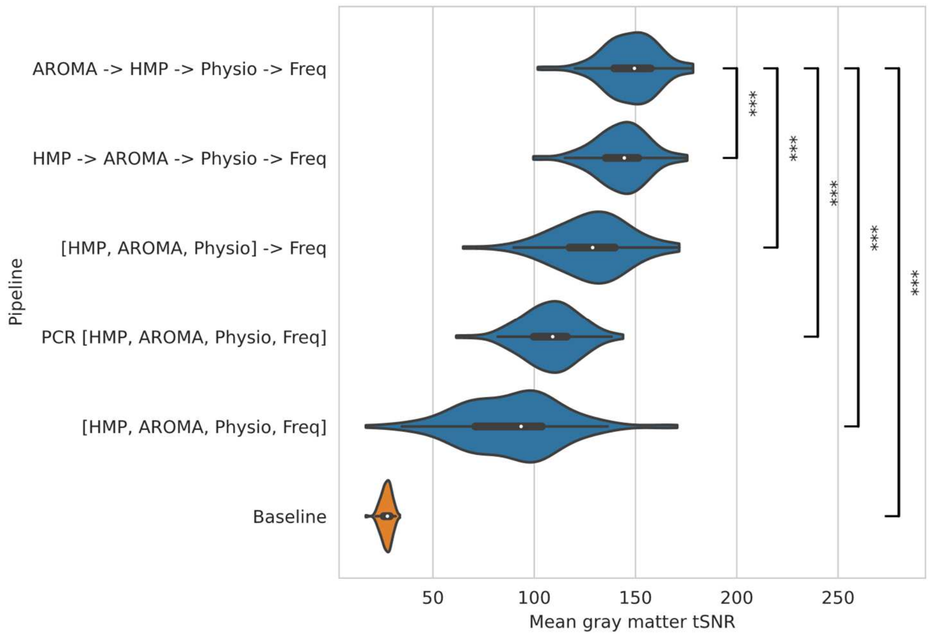


**Figure S16**. Temporal signal-to-noise ratio (tSNR) compared across HCP images processed with each pipeline. tSNR was computed for each voxel as the mean signal divided by the temporal standard deviation, then averaged across gray matter voxels. Assuming the temporal standard deviation is mainly attributable to motion rather than neuronal activation, higher tSNR is preferred.
